# Selection on growth rate and local adaptation drive genomic adaptation during experimental range expansions in the protist *Tetrahymena thermophila*

**DOI:** 10.1101/2021.02.04.429706

**Authors:** Felix Moerman, Emanuel A. Fronhofer, Florian Altermatt, Andreas Wagner

**Affiliations:** Department of Evolutionary Biology and Environmental Studies, University of Zürich, Winterthurerstrasse 190, Zürich CH-8057, Switzerland; Department of Aquatic Ecology, Eawag: Swiss Federal Institute of Aquatic Science and Technology, Überlandstrasse 133, Dübendorf CH-8600, Switzerland; Swiss Institute of Bioinformatics, Quartier Sorge—Bâtiment Génopode, Lausanne 1015, Switzerland; ISEM, Univ Montpellier, CNRS, EPHE, IRD, Montpellier, France; The Santa Fe Institute, Santa Fe, New Mexico 87501, USA; Stellenbosch Institute for Advanced Study (STIAS), Wallenberg Research Centre at Stellenbosch University, Stellenbosch 7600, South Africa

**Keywords:** gene ontology, life-history, pH, range expansions, whole genome resequencing, Tetrahymena

## Abstract

1. Populations that expand their range can undergo rapid evolutionary adaptation of life-history traits, dispersal behaviour, and adaptation to the local environment. Such adaptation may be aided or hindered by sexual reproduction, depending on the context.
2. However, few studies have investigated the genomic causes and consequences or genetic architecture of such adaptation during range expansions.
3. We here studied genomic adaptation during experimental range expansions of the protist *Tetrahymena thermophila* in landscapes with a uniform environment or a pH-gradient. Specifically, we investigated two aspects of genomic adaptation during range expansion. Firstly, we investigated the genetic architecture of adaptation in terms of the underlying numbers of allele frequency changes from standing genetic variation and *de novo* variants. We focused on how sexual reproduction may alter this genetic architecture. Secondly, identified genes subject to selection caused by the expanding range itself, and directional selection due to the presence or absence of the pH-gradient. We focused this analysis on alleles with large frequency changes that occurred in parallel in more than one population to identify the most likely candidate targets of selection.
4. We found that sexual reproduction altered genetic architecture both in terms of *de novo* variants and standing genetic variation. However, sexual reproduction affected allele frequency changes in standing genetic variation only in the absence of long-distance gene flow. Adaptation to the range expansion affected genes involved in cell divisions and DNA repair, whereas adaptation to the pH-gradient additionally affected genes involved in ion balance, and oxidoreductase reactions. These genetic changes may result from selection on growth and adaptation to low pH.
5. Our results suggest that the evolution of life-history and the adaptation to the local environment has a genetic basis during our range expansion experiment. In the absence of gene flow, sexual reproduction may have aided genetic adaptation. Gene flow may have swamped expanding populations with maladapted alleles, thus reducing the extent of evolutionary adaptation during range expansion. Sexual reproduction also altered the genetic architecture of our evolving populations via *de novo* variants, possibly by purging deleterious mutations or by revealing fitness benefits of rare genetic variants.

## Introduction

Range expansions and biological invasions are increasingly common in many species. They often result from anthropogenic disturbances or species introductions (Parmesan et al., 1999; Kowarik, 2003; Chen et al., 2011). Because range expansions can have dramatic consequences on species distributions and ecosystems (see for example Grosholz, 2002; Phillips et al., 2009), it is important to understand their ecological and evolutionary consequences. Theory predicts that expanding populations can undergo adaptation through two major mechanisms. Firstly, spatial sorting of individuals during range expansions is expected to lead to increased dispersal behaviour at the range edge (Shine et al., 2011; Phillips and Perkins, 2019). Secondly, natural selection at the low population density range edge is expected to lead to selection for an r-selected life-history strategy (Phillips, 2009; Burton et al., 2010; Phillips et al., 2010). Recent studies have demonstrated that expanding populations can indeed rapidly evolve their dispersal behaviour (Brown et al., 2007; Williams et al., 2016; Ochocki and Miller, 2017) and life-history strategy (Phillips, 2009; Fronhofer and Altermatt, 2015). Additionally, they can evolve specific adaptations to local abiotic conditions, such as temperature and novel food sources (Van Petegem et al., 2016; Szűcs et al., 2017). Because selection pressures during range expansion can be strong, populations at a range edge may adapt quickly (Sakai et al., 2001). However, evolution during range expansion does not necessarily affect both dispersal behaviour and life-history traits, especially when these traits trade-off with each other (Burton et al., 2010; Clarke et al., 2019).

A population’s potential to adapt evolutionarily may be constrained if the adapting traits have a complex genetic architecture (Mackay, 2001; Hansen, 2006). Such an architecture may exist for traits such as growth rate, dispersal behaviour, and adaptation (Stearns, 2000; Hansen, 2006; Saastamoinen et al., 2018). Multiple traits with a complex genetic architecture can further reduce evolutionary potential. One important mechanism that can help overcome evolutionary constraints imposed by genetic architecture is the reshuffling of existing genetic variation through sexual reproduction (Bell, 1982; Maynard-Smith, 1978; Otto and Lenormand, 2002). This benefit of sexual reproduction has been demonstrated repeatedly (Colegrave, 2002; McDonald et al., 2016; Luijckx et al., 2017; Petkovic and Colegrave, 2019), and is especially pronounced in the presence of high genetic diversity (Lachapelle and Bell, 2012). Increased genetic variation or an influx of locally adapted genes can also increase the success of invasions and range expansions (Schmeller et al., 2005; Lavergne and Molofsky, 2007; Currat et al., 2008). However, sexual reproduction during range expansions can also hinder adaptation, for example in the process of ”gene swamping”. Gene swamping occurs when populations expand their range against an environmental gradient, and when genes ill-adapted to the range edge flow from a high-density core population to a low-density population at the range edge. In asexual populations, such maladapted individuals may simply disappear, as they fail to survive. In contrast sexual populations may experience a general decline in fitness, when enough individuals move to the range core, so that all offspring descend from maladapted individuals. According to the gene swamping hypothesis, gene flow can decelerate adaptation of a population at the range edge (Haldane and Ford, 1956; García-Ramos and Kirkpatrick, 1997; Kirkpatrick and Barton, 1997; Polechová and Barton, 2015; Polechová, 2018), as recently demonstrated both in an experimental study (Moerman et al., 2020b) and a field study (Bachmann et al., 2020).

However, little is known of the genetic mechanisms behind range expansions. With few exceptions (Bosshard et al., 2020), studies on the evolutionary genetics of range expansions have focused on genetic population structure and diversity (see for example Swaegers et al., 2013; Bors et al., 2019). This work has demonstrated that, first, populations at the range edge typically harbour less genetic variation than the core populations (Excoffier et al., 2009). Second, genetic drift can strongly affect populations at a range edge, which can lead to ”allele surfing”, the increase in frequency and potential fixation of neutral or maladaptive variants at a range edge (Klopfstein et al., 2006; Excoffier et al., 2009). Both these mechanisms can constrain the adaptive potential at a range edge. Despite such work, the genetic basis of phenotypic evolution during range expansions remains poorly understood. For example, we know little about the kinds of genes that are involved in phenotypic evolution during range expansion. We know even less about how gene swamping may alter the genetic architecture of adaptation during range expansion.

We here studied the population genomics of previously published range expansion experiment with the protist *Tetrahymena thermophila* (Moerman et al., 2020b). In this experiment, we had experimentally investigated the gene swamping hypothesis, by assessing how sexual reproduction and gene flow affected evolution during range expansion either into a uniform environment or into a pH-gradient. We chose a pH-gradient for this experiment for two reasons. Firstly, pH is an important environmental stressor linked to ocean acidification (Caldeira and Wickett, 2003; Raven et al., 2005; Zeebe et al., 2008). Secondly, pH can easily be manipulated experimentally. The experiment showed that sexual reproduction aided adaptation during range expansion, but only in the absence of gene flow. We here performed whole genome sequencing of pooled *T. thermophila* populations from this experiment. We identified the populations with the strongest phenotypic changes, pooled individuals from these populations, and sequenced the genomic DNA of the resulting pools. We then investigated how genetic architecture, in terms of genetic changes from standing variation and from *de novo* variants, was affected by sexual reproduction, gene flow, and a pH-gradient. Genetic architecture describes the genetic effects that affect a phenotypic character and can include allelic changes and mutations, pleiotropy, dominance and epistatic interactions (Hansen, 2006). We focus in this study on allelic changes from standing variation and *de novo* variants. We predicted that the amount of genetic change reflects the amount of phenotypic change, and that sexual reproduction could facilitate genetic change by bringing together adaptive variants, but only in the absence of gene swamping. Furthermore, we identified genes involved in adaptation to the range expansion itself and to the pH-gradient, focusing on selective sweeps of large effect alleles that occurred in parallel in multiple populations. Here we predicted that adaptation during range expansion may act on life-history traits or dispersal behaviour, whereas adaptation to the pH-gradient may act predominantly on the ability to cope with acidity, either through ion maintenance or cell membrane function.

## Material and methods

### Study organism and experimental evolution

*Tetrahymena thermophila* is a freshwater ciliate commonly used in ecological and evolutionary experiments (Collins, 2012; Altermatt et al., 2015; Cairns et al., 2020; Moerman et al., 2020b), including in studies of range expansions (Giometto et al., 2014; Fronhofer and Altermatt, 2015; Giometto et al., 2017). *Tetrahymena* is characterized by nuclear dualism. That is, cells carry a polyploid (n=45) macronucleus and a diploid micronucleus (Lynn and Doerder, 2012). *Tetrahymena* usually reproduces asexually, but can be induced to reproduce sexually through starvation (Cassidy-Hanley, 2012). During asexual reproduction, both nuclei are copied, but the chromosome copies of the macronucleus are partitioned randomly among daughter cells (Ruehle et al., 2016). We used ancestral and evolved *Tetrahymena* populations from a range expansion experiment by Moerman et al. (2020b) to investigate the genomic basis of adaptation during range expansions. We explain the most pertinent details of this experiment here and refer the reader for other details to Moerman et al. (2020b).

Briefly, in this experiment, we mixed four phenotypically divergent (Moerman et al., 2020a) clonal strains of *T. thermophila* to create a genetically diverse ancestral population. For details on strain identities and the mixing of the ancestral population, see section S1.1 in the Supplementary Material. Subsequently, we allowed 40 replicates of this ancestral population to expand their range for ten weeks (250 generations). We kept all stock cultures and populations during experimental evolution in a climate room at 20 °C.

During experimental range expansion, we used an established system of two-patch landscapes (Fronhofer and Altermatt, 2015) to emulate an expanding range edge. These two-patch landscapes consist of two Sartedt tubes (25 mL) that contain 15 mL of modified Neff medium (Cassidy-Hanley, 2012), and are connected by an eight cm silicone tube. A plastic clamp is used to close this tube. When open, the tube allows cells to actively disperse between the two patches (see figure 1). To emulate the range core, we maintained the ancestor clones in separate glass tubes, in which they experienced continuously high population densities and slow division rates. These conditions experienced by the ”range core” populations are identical to the conditions experienced by these clones while being maintained in the laboratory. We initiated the experiment by inoculating one patch of the two-patch landscape (the ”home patch”) with 200 µL of the ancestral population. During range expansion, we controlled three factors. Firstly, we controlled the abiotic environment, where populations either experienced a ”uniform” environment, with a pH equal to 6.5, or a ”pH-gradient” environment, where the pH gradually decreased from 6.5 of 4.0 during range expansion, to emulate an environmental pH-gradient. We determined this pH range as the pH of fresh Neff-medium (pH 6.5), and the lowest pH allowing cells to grow (pH 4.0). Secondly, we controlled the reproductive mode, with ”asexual” populations experiencing only asexual reproduction during the experiment, whereas ”sexual” populations also reproduced mainly asexually, but experienced four sexual reproduction events during the experiment. Lastly, we controlled long-distance gene flow from the range core to the edge, where either long distance gene flow was ”absent”, meaning we didn’t impose long-distance gene flow, or gene flow was ”present”, meaning that we imposed four long-distance gene flow events, by replacing part of the population at the range edge with individuals from the range core. Altogether, our experiment thus used eight treatments. For each treatment, we evolved five replicate populations, resulting in a total of 40 populations.

**Figure 1:**
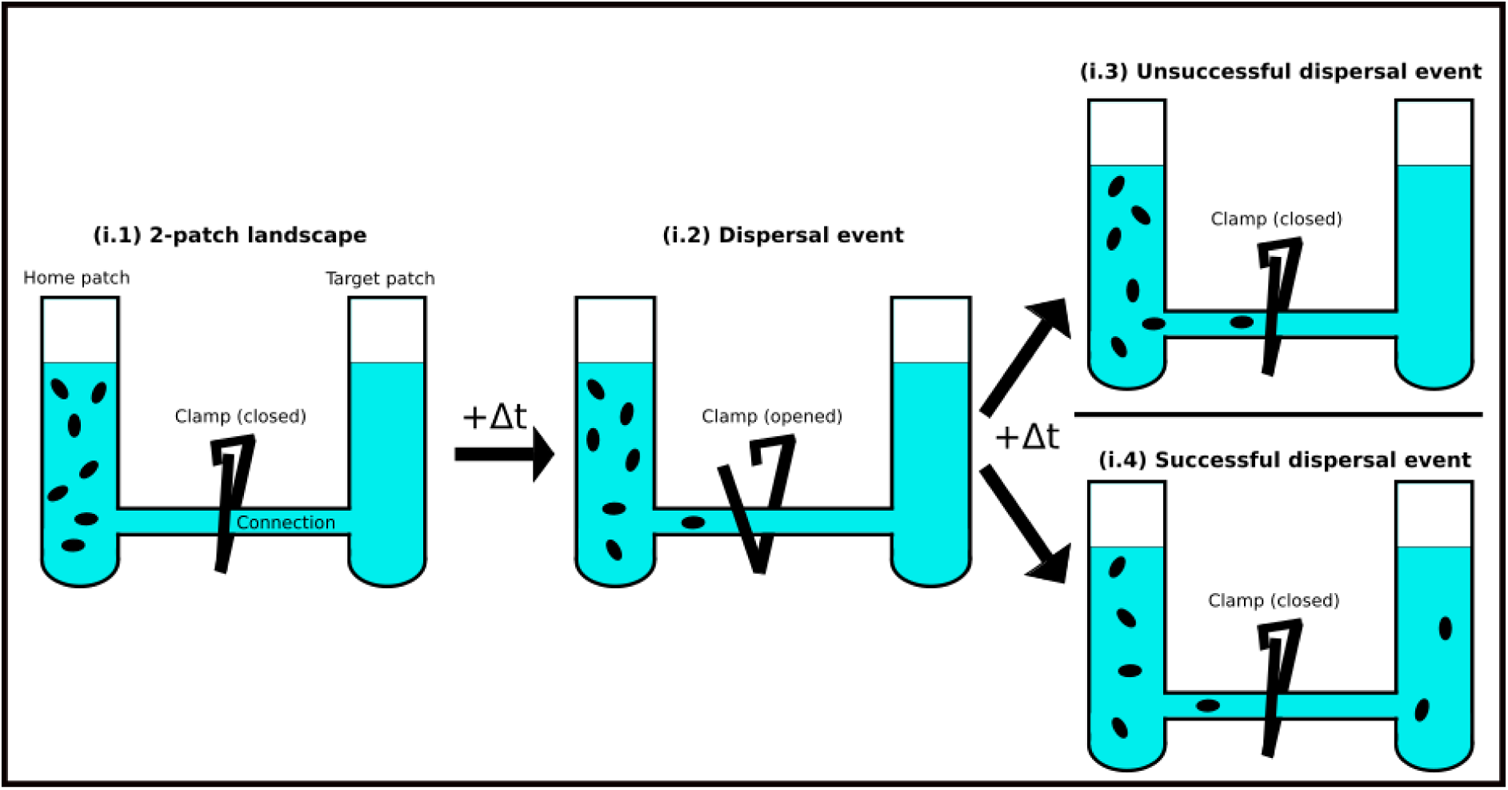
Two-patch landscapes. i.1: Two-patch landscapes consist of connected tubes. i.2: To initiate dispersal, the connection is opened for one hour, allowing cells to actively swim from the home patch to the target patch. i.3: If we found no cells in the target patch, dispersal was deemed unsuccessful. i.4: If we did detect cells in the target patch, dispersal was deemed successful.

We then subjected the populations to ten weeks of experimental evolution, during which we repeated the same cyclical procedure every 14 days. Specifically, we subjected each population to three dispersal events, which took place on days 1, 3 and 5 of the 14-day cycle. After the third dispersal event, we subjected populations of the appropriate treatment groups to a long-distance gene flow and/or sexual reproduction event on day 8. Lastly, we subjected the populations to two additional dispersal events on days 10 and 12. Consequently, our experiment included four distinct gene flow and sexual reproduction events, in week two, four, six and eight of the experiment. Over the entire experiment, population densities remained high (10^4^ − 10^6^ individuals). During the dispersal events, between 1 % and 20 % of our populations typically dispersed, such that dispersal never led to extreme bottlenecks. Therefore, genetic drift is likely negligible on the timescale of this experiment (Hartl and Clark, 2006). In consequence, genetic changes are likely due to adaptive evolution, or hitchhiking. For a detailed description and a visual representation of the experimental design, see the Supplementary Material sections S1.3 and S1.4.

### Common garden and growth assessment

After a common garden phase of 18 generations to reduce epigenetic and maternal effects, we assessed the fitness change of the evolved populations compared to the ancestral population. To do so, we performed growth assays for the evolved and ancestral populations under multiple pH-values (pH 6.5, 6.0, 5.5, 5.0, 4.5, 4.0, 3.5 and 3.0). We then fit a Beverton-Holt population growth model (Beverton and Holt, 1993) to infer the intrinsic population growth rate *r*_0_ (see Supplementary Material section S1.5). We then calculated the change in fitness that occurred in each evolved population as the change in the population’s intrinsic growth rate *r*_0_ relative to the ancestral population. More specifically, we calculated the log-ratio response as

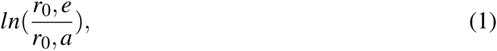

where *r*_0_*, e* and *r*_0_*, a* are the intrinsic population growth rate of the evolved population and the ancestral population, respectively.

### Sequencing and identification of variants

To identify candidate loci for genetic adaptation, we performed population level sequencing (”PoolSeq”; Schlötterer et al., 2014).

We only wished to sequence the macronuclear DNA, because only the expressed macronuclear DNA affects the phenotype. We thus first separated macronuclei from micronuclei. Following nuclear separation, we extracted DNA from each population. To obtain statistically sound results the recommended sequencing depth for whole genome population resequencing is between 50X and 200X (Schlötterer et al., 2014; Kofler and Schlötterer, 2014). Because of the large genome size of *T. thermophila* (105 Mb; Eisen et al., 2006), it was only possible to sequence a limited number of populations at this depth. We therefore selected two populations for genomic sequencing from each treatment group, based on the fitness change these populations experienced. That is, we sequenced the populations with the largest increase in fitness, because they were the most likely candidates to show adaptive genetic changes. In this way, we obtained sequencing data for 16 (=2 × 8 treatments) populations. Next, we mapped the obtained sequences to the *T. thermophila* reference genome, and performed variant calling to identify genetic variants. In analyzing the resulting data, we studied standing genetic variation and its allele frequency change, as well as variants that had occurred *de novo*. In the next sections, we describe each of these steps in detail.

#### Nuclear separation and DNA extraction

We separated macronuclei and micronuclei of the ancestral clones and of all 34 surviving evolved populations using a differential centrifugation protocol adapted from Sweet and Allis (2006, see Supplementary Material section S2 for details). We obtained for every population three pellets containing macronuclei (pellet P1, P2 and P3). We extracted DNA from each of the pellets using a Qiagen^®^ DNeasy Blood and Tissue kit (Cat No./ID: 69506), following the manufacturer’s protocol (see https://www.qiagen.com/us/resources/resourcedetail?id=63e22fd7-6eed-4bcb-8097-7ec77bcd4de6&lang=en). After DNA extraction, we determined the quality and quantity of the DNA using a Nanodrop spectrophotometer (model ND-1000) and selected for each population the pellet with the highest quality (see Supplementary Material section S3). We stored DNA samples in a −80 °C freezer until sequencing. Library preparation and sequencing was performed by the Functional Genomics Center Zürich, which prepared 2×150 bp paired-end libraries and sequenced them using the Illumina Novaseq 6000 platform. We aimed to obtain population level DNA-sequences at approximately 100x genome coverage (see Supplementary Material section S4.2 for coverage statistics).

#### Bioinformatic pipeline

We trimmed the reads of all fastq files using Trimmomatic version 0.39 (Bolger et al., 2014) to remove Illumina adapter sequences (ILLUMINACLIP option) and low quality segments (SLIDINGWINDOW:4:15). Additionally, we removed any reads falling below a length of 36 bp (MINLEN:36). After trimming, we mapped the reads for each of the 20 populations to the *Tetrahymena thermophila* macronuclear reference genome (Eisen et al., 2006) using the Burrows-Wheeler aligner (Li and Durbin, 2009) version 0.7.17-r1188. On average, 97.7% of the reads aligned to the reference genome (see Supplementary Material section S4.1 for mapping rates of all populations).

We then called genetic variants using the BCFtools multiallelic caller (Danecek et al., 2016). We first called variants to identify sites where the four ancestral and 16 evolved populations differed from the reference genome, and stored the resulting allele counts.

We mapped all variants to genes using bedtools (v2.27.1; Quinlan and Hall, 2010). To this end, we first obtained the genome annotation file from the *T. thermophila* reference genome (Eisen et al., 2006), and filtered this file to only keep entries corresponding to protein-coding genes. We then used the intersect function in bedtools to map the variants to the gene entries.

#### Quantifying allele frequencies

To detect changes in allele frequencies, we first calculated the frequency of each allele present in the ancestral population (expected allele frequency). Because we had sequenced the ancestor clones individually, we started by identifying all positions along the genome where at least one of the ancestral clones differed from the reference genome. We then calculated for these positions the allele frequency in the ancestral population (see Supplementary Material section S5). Subsequently, we filtered these positions to exclude positions with low sequencing quality (see Supplementary Material section S6.1 for details). We then calculated the allele frequencies for the 16 evolved populations at all positions along the genome that had not been eliminated by filtering. For each of the evolved populations, this resulted in a list of positions where genetic variation had initially been present in the ancestral population, and where allele frequency changes were therefore possible.

#### Identifying *de novo* variants

To detect *de novo* variants, we kept only those variants that differed from the reference genome in an evolved population, but not so in any of the four ancestral clones. We then removed any positions with low sequencing quality (see Supplementary Material section S6.2 for details).

### Statistical analyses

We performed statistical analyses, unless otherwise specified, using the R statistical language version 4.0.2 (R Core Team, 2020).

#### Analyzing fitness changes

We applied the fitness assay described in Moerman et al. (2020b) to the populations included in the genomic analysis. Briefly, we determined the change in the intrinsic population growth rate *r*_0_ of each evolved population relative to the ancestral population in the pH conditions experienced during range expansion (pH 6.5 for populations expanding into a uniform environment, and pH 4.0 for populations expanding into a pH-gradient). For details, see section S11 in the Supplementary Material.

#### Quantifying frequency changes in standing genetic variation

We determined for all polymorphic sites in the ancestral population if the alleles at the site had experienced a significant change in frequency. To do so, we performed a binomial test comparing the number of reference and non-reference reads in evolved populations with the expected allele frequency of the ancestral population. Because we aimed at identifying the most likely candidates of selection, we used a p-value of 0.001 in this test, to which we applied a Bonferroni correction to avoid falsely detecting allele frequency changes due to multiple testing. We therefore considered only variants with a p-value smaller than 10^−10^ as significantly different from the expectation. Such a stringent cut-off implies a low false positive rate is low, but it also implies that we may miss some alleles whose frequency changed due to selection (Kofler and Schlötterer, 2014).

In a preliminary analysis of standing genetic variation, we observed that the ancestor clone SB3539 showed a number of anomalies that made its exclusion from further analysis desirable. First, this clone contained approximately 30 times more non-reference alleles than the other ancestral clones and than each of the evolved populations (Table S7). Second, clone SB3539 deviated strongly in its transition/transversion ratio (clone SB3539: 0.9344; other populations 0.6544—0.7611; Table S7). Furthermore, selection acted very strongly against this clone as a whole, as evidenced by post-evolution allele frequencies that were dramatically different from those in all other populations (Fig. S4). Relatedly, all evolved populations showed an extremely high number of variants (approx. 1.1 × 10^4^) that changed in their frequency by approximately 0.25, a value that corresponds closely to the initial frequency of clone SB3539 (27% of the ancestral population; Table S1). To avoid potential confounding effects due to this clonal selection, we limited our analyses to those alleles whose frequency changed by an amount that is larger than could possibly be caused by selection against this clone. Specifically, we only analyzed alleles whose frequency changed by a value of 0.3 or higher (rounded up from 0.27, the starting frequency of clone SB3539).

Next, we assessed if the number of alleles whose frequency changed significantly was affected by reproductive mode, gene flow, and the pH-gradient. To do so, we counted for every evolved population the number of variants that significantly changed in frequency. In this analysis, we applied various cut-offs (0.3, 0.4, 0.5, 0.6, 0.7 and 0.8) for the magnitude of the allele frequency change, i.e., the minimum absolute difference between the expected and the observed allele frequency, required for variants to be included. We fit a Bayesian generalized linear mixed model with a Poisson distribution to the resulting data, using the “Rstan” and “statistical rethinking” packages (Stan Development Team, 2020; McElreath, 2015). We created a model in which reproduction, gene flow, abiotic conditions and the cut-off value are all allowed to interact as fixed effects, and used the identity of each replicate evolved population as a random effect. We report posterior distributions (means and 95 % confidence intervals) for the parameter estimates. Additionally, we plotted the population’s fitness change against the number of alleles that changed significantly in frequency. We calculated the Pearson correlation between the two metrics (paired sample correlation test), and did so for different cut-off values of the magnitude of allele frequency change.

#### Quantifying *de novo* variants

To assess how reproductive mode, gene flow, and the pH-gradient affect the evolutionary dynamics of *de novo* variants in our populations, we counted the number of novel variants in each of the evolved populations. To this end, we used eight cut-off values of increasing allele frequencies, ranging from 0.1 to 0.9. For each of these values we counted the number of *de novo* variants that had reached an allele frequency larger than the cut-off. We then fit a Bayesian linear mixed model with Rstan and the statistical rethinking package (Stan Development Team, 2020; McElreath, 2015) using reproduction, gene flow, presence of a pH-gradient, and cut-off as fixed effects, and replicate population as a random effect. We report posterior distributions (means and 95 % confidence intervals) for the parameter estimates.

#### General adaptations during range expansion

To identify allele frequency changes due to general adaptation — they were subject to selection regardless of reproductive mode, gene flow, or the pH-gradient — we focused on alleles whose frequency increased or decreased consistently across all or most populations. Specifically, we identified allelic variants that were present in the ancestral population, that changed their frequency not merely as a result of the global selection we observed against clone SB3539 (i.e., they changed their frequency by more than 0.3), and that changed significantly in frequency (p-value<10^−10^). Among the remaining alleles, we then considered only those whose frequency changed in the same direction for 75% (12/16) of all evolved populations. We call the resulting dataset the ”general adaptation dataset”.

#### Gradient-specific adaptations: Cochran-Mantel-Haenszel test

To identify directional allelic changes due to the presence/absence of the pH-gradient, we performed a Cochran-Mantel-Haenszel test (PoPoolation2-package; Kofler et al., 2011). This test compares pairs of populations and it can identify genomic loci that show consistently different allele frequencies. To account for the effect of reproduction and gene flow, we always compared pairs of populations with the same reproduction or gene flow treatment. For example, we compared an asexual population without gene flow expanding into a pH-gradient with another asexual population without gene flow but expanding into a uniform environment (see Supplementary Material section S8.1). With this approach, we aimed at detecting loci that display consistent differences between populations expanding into a pH-gradient and populations expanding into a uniform environment, independent of gene flow and reproductive mode. To account for multiple testing, we applied a Bonferroni correction (Bland and Altman, 1995). Because our genomes harboured approximately 10^5^ genetically variable positions, and we wished to use a conservative p-value of 0.001, we only classified positions with a p-value smaller than 10^−8^ as significant. We refer to this dataset as the ”gradient-specific adaptation” dataset. We again focused on alleles within protein-coding genes and examined the functional annotations of these genes.

#### Gene ontology enrichment

To assess whether allele frequency changes occur in genes associated with specific functions, we performed a gene ontology (GO) term enrichment analysis using GOWINDA (version 1.12; Kofler and Schlötterer, 2012). This analysis identifies terms describing the molecular or biological function and cellular component associated with each gene, and then assesses which of such terms are over-represented in a dataset. We first queried the Uniprot Knowledgebase to obtain all entries associated with the *T. thermophila* reference genome (TaxID: 312017). This resulted in a list of genes encoded in the Tetrahymena genome together with GO-terms associated with each gene. We then performed a GOWINDA analysis using this dataset as a reference database, to identify which gene ontology terms were significantly enriched. We performed this analysis for two sets of genes, the genes associated with general adaptations and the genes associated with gradient-specific adaptations.

For both analyses, we kept only those GO-terms with an FDR (false discovery rate) value below 0.05, following a Benjamini-Hochberg correction (Benjamini and Hochberg, 1995). Next, we grouped the enriched GO-terms into 13 categories of cellular mechanisms (see Supplementary Material table S15). We then compared the proportion of all GO-terms associated with the 13 categories between general adaptation genes and gradient-specific genes, using a *χ*^2^-test. Finally, we performed a post-hoc test to assess which categories are differently enriched in the gradient-specific genes and the general adaptation genes (Ebbert, 2019).

## Results and discussion

### Genetic architecture of adaptation

By reshuffling genetic variation, sexual reproduction can have important consequences for adaptive evolution (Maynard-Smith, 1978; Bell, 1982; Otto, 2009). This reshuffling of genetic variation may affect both existing variation and new variants. To investigate how gene swamping may affect the genetic architecture during range expansions, we here assessed how sexual reproduction altered phenotypic change and genetic change through standing variation and *de novo* variants, in landscapes with/without a pH-gradient and in presence/absence of gene flow.

#### Gene swamping affects fitness change

We repeated the analysis described in Moerman et al. (2020b) using only those 16 populations that we included in the genetic analysis, to ensure that the results remain unchanged.

We observed a positive change in fitness for all populations included in the genetic analysis (Figure 2). The fitness increase was stronger for populations that expanded into a pH-gradient (panels B and D in Figure 2) than for populations that expanded into a uniform environment (Figure 2 panels A and C; *χ*^2^_1,16_=299.16, p<0.001). In both environments (pH-gradient/uniform), the presence of gene flow had a positive effect on fitness (*χ*^2^_1,16_=8.09, p=0.016), as did sexual reproduction (*χ*^2^_1,16_=6.44, p=0.027). However, there was a significant interaction between sexual reproduction and the presence of gene flow (*χ*^2^_1,16_=14.36, p=0.003), suggesting that sexual reproduction only provided benefits if gene flow was absent. The results are congruent with those described in Moerman et al. (2020b).

**Figure 2:**
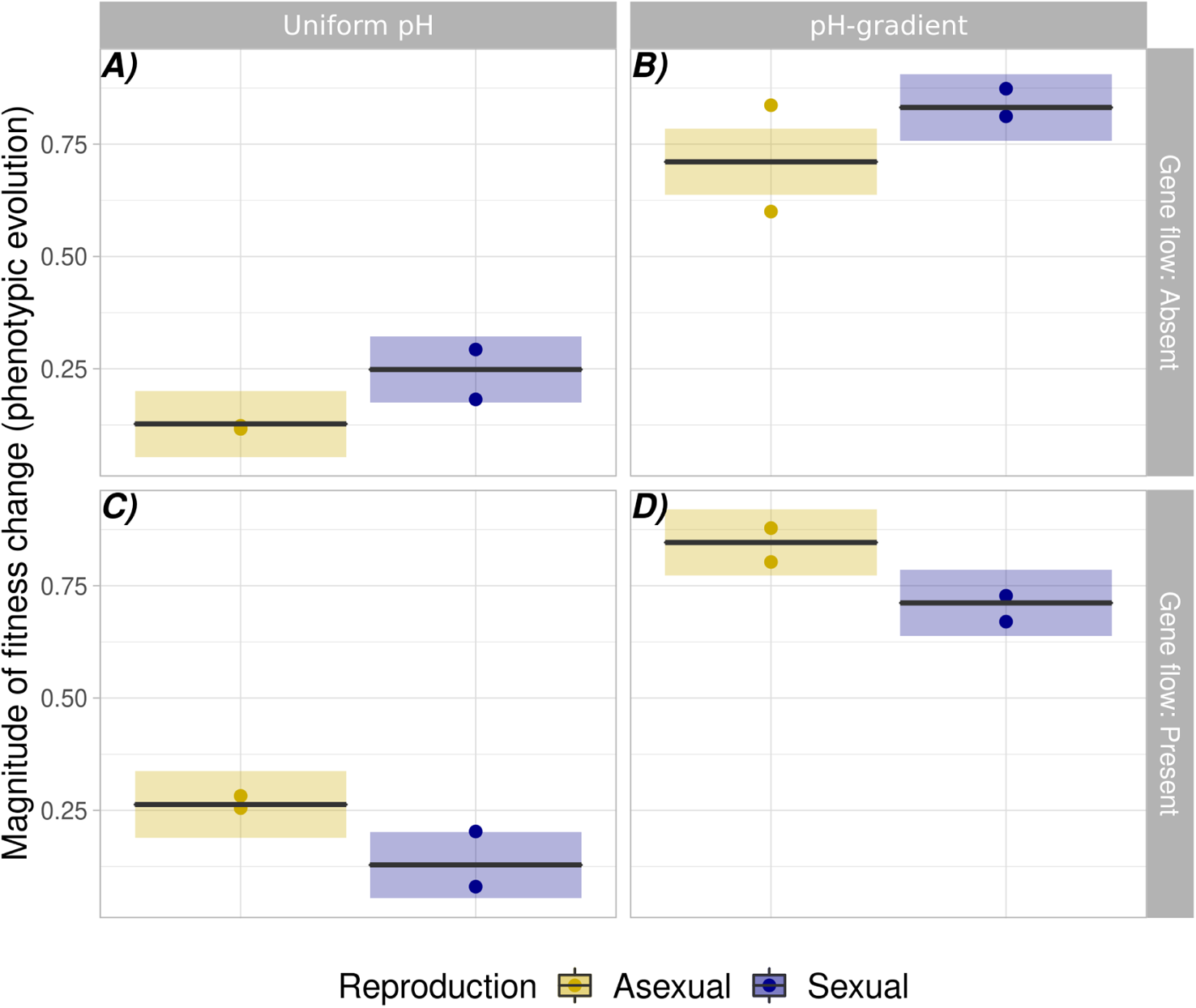
Gene swamping affects fitness changes during range expansion. Fitness change expressed as the log-ratio change in intrinsic population growth rate, calculated as the ratio of the logarithm (base 2) of the intrinsic population growth rate (*r*_0_) of an evolved population and the intrinsic population growth rate (*r*_0_) of the ancestral population. Circles represent data for individual populations. Black lines and shaded areas represent mean predictions and 95% confidence intervals of the best model. Colours represent mode of reproduction (yellow=asexual, blue=sexual). Left subplots (panels A and C) show data and model predictions for populations expanding into a uniform environment, right subplots (panels B and D) for populations expanding into a pH-gradient. Upper subplots (panels A and B) show data and model predictions for populations where gene flow is absent, and lower subplots (panel C and D) for populations where gene flow is present. Adapted from Moerman et al. (2020b).

#### Frequencies of *de novo* variants change most strongly in sexually reproducing populations

In a second analysis, we investigated the presence of *de novo* variants that rose to detectable allele frequencies. We found a total of 27,766 *de novo* variants across all 16 evolved populations. They reached frequencies between 10% and 100%. Only 0.02% of these *de novo* variants went to fixation. We found that 31.73 % of *de novo* variants occurred in protein-coding genes. Given that 47.87 % of the *Tetrahymena* genome is protein-coding (Eisen et al., 2006), *de novo* variants are less likely to occur in protein coding regions than expected by chance (exact binomial test, p <0.001). Any one non-coding *de novo* variants may have hitch-hiked to high allele frequency due to its physical proximity to an adaptive coding variant (Charlesworth et al., 2000). Alternatively, it may itself have been subject to positive selection, for example by altering DNA binding sites of transcription factors and thus gene regulation in a beneficial way (Latchman, 1997; Spitz and Furlong, 2012). Most (82.30%) *de novo* variants were indels. Significantly fewer indels (29.14%) than SNPs (46.36%) occurred in protein-coding regions (Pearson Chisquared test: *χ*^2^_1_=130.89; p <0.001). Only *de novo* indels but not SNPs were less likely to occur in protein-coding regions than expected by chance (indels: 29.14%; p <0.001; SNPs; 46.36%, p=0.066, exact binomial test). These observations are consistent with the notion that indels inside protein-coding regions are more likely to be deleterious than SNPs, because they are more likely to disrupt open reading frames.

We then studied the role of reproductive mode, gene flow, and the pH-gradient as explanatory variables for the number of *de novo* variants (response variable). We detected similar numbers of *de novo* variants in populations expanding into a uniform environment (Figure 3, panels A and C) and into a pH-gradient (Figure 3, panel B and D). We observed more variants in sexually reproducing populations (Figure 3, blue lines and circles) than in asexual populations (Figure 3, yellow lines and circles), but only at low values for the cut-off (Reproduction×Cutoff). See Supplementary Material section S13 for full posterior distributions.

**Figure 3:**
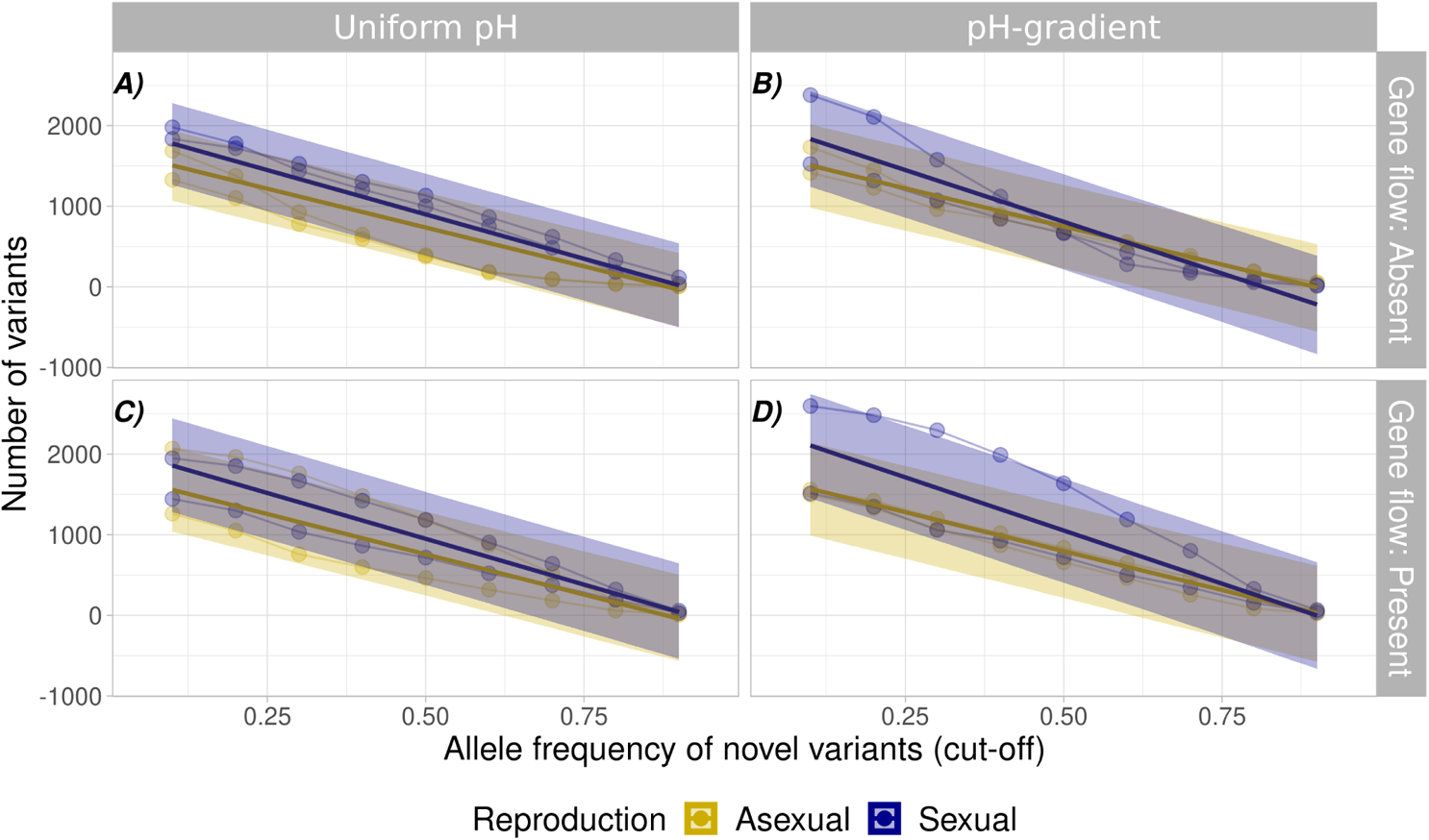
*De novo* variants increase to higher frequency in sexual populations. The figure shows the number of novel variants for the evolved populations expanding into a uniform environment (panels A and C) or into a pH-gradient (panel B and D). The x-axis shows the cut-off value in minimum allele frequency. The y-axis shows the number of *de novo* variants whose frequency is above a given cut-off. Circles represent the data, with thin lines connecting data from the same replicate population. The thicker opaque lines and shaded areas show the predicted means and 95% confidence intervals for the best model. Colours represent the mode of reproduction.

Because most new mutations are neutral or deleterious (Loewe and Hill, 2010), the accumulation of mutations in a population often leads to a decrease in fitness. Sexual reproduction can help avert this decrease by purging deleterious mutations and separating them from rare beneficial mutations (Muller, 1932; Maynard-Smith, 1978; Peck, 1994; Fisher, 2000; Johnson and Barton, 2002; McDonald et al., 2016). This may explain why more *de novo* variants rose to detectable allele frequency in sexually reproducing populations. Because the chromosomes of the polyploid macronucleus are partitioned randomly among the offspring of asexually reproducing *T. thermophila*, different offspring of the same individual may vary in the copy numbers of specific variants (Ruehle et al., 2016). Therefore, in asexual *T. thermophila* populations, selection likely acts against individuals carrying deleterious variants at a high copy number. In contrast, sexual recombination may reduce this mutational load by separating maladaptive mutations from neutral or beneficial mutations (Maynard-Smith, 1978). Given the large number of variants we identified, an alternative explanation could be that some genetic variants we had classified as *de novo* were actually present at the start of our experiment, but at an undetectably low frequency. As discussed in Hansen (2006), some such variants may be beneficial, but these benefits may be hidden, for example if a variant co-occurs with a deleterious variant that offsets its advantage. Such benefits can be revealed by sexual reproduction. Therefore, some initially very rare variants may have risen to high frequency in our populations following sexual reproduction.

#### Reproduction alters allele frequency change when gene flow is absent

Next, we assessed how reproductive mode, gene flow and the presence of a pH-gradient affected standing genetic variation in our evolving populations. After correcting for multiple testing and clonal selection on one of our ancestral clones (see Methods, ”Statistical analyses”), we observed significant allele frequency changes in 13,278 alleles. Only 29.39% of alleles occurred in protein-coding regions of the genome. The percentage of macronuclear DNA predicted to occur in protein-coding genes of *T. thermophila* equals 47.78% (Eisen et al., 2006). This means that we observed significantly fewer allele frequency changes in protein-coding genes than expected by chance alone (exact binomial test: p <0.001). The abundance of non-coding alleles with significant frequency changes is consistent with the possibility that gene regulatory changes are important for evolutionary adaptation in our populations. Alternatively, non-coding variants may frequently hitch-hike to high frequency with protein coding variants. More than half (57.1%) of all alleles were indels as opposed to single nucleotide polymorphisms (42.9%). More indels and SNPs than expected by chance occurred in non-coding regions (exact binomial test; p <0.001 for both). However, there were significantly more SNPs (34.61%) than indels (25.46%; Pearson Chi-squared test: *χ*^2^_1_=130.89; p <0.001) in protein-coding regions of the genome, consistent with the expectation that indels are likely to be more deleterious than SNPs.

We then fit a Bayesian generalized linear mixed model that included the mode of reproduction, gene flow, and abiotic conditions as explanatory variables, the number of alleles which changed in frequency as the response variable, and the identity of the evolved population as a random effect.

The total number of alleles that changed significantly in frequency was similar for populations expanding into a uniform environment (Figure 4, panels A and C) and into a pH-gradient (Figure 4, panels B and D). However, the effect of reproduction and gene flow differed between these populations. In the absence of gene flow, we observed more genetic changes for populations with a sexual mode of reproduction when populations expanded into a uniform environment (Figure 4, panel A), but fewer genetic changes when populations expanded into a pH-gradient (Figure 4, panel B). When populations experienced gene flow during range expansion (Figure 4, panels C and D), there was no clear difference between asexual and sexual populations. Full posterior distributions can be found in Supplementary Material section S12.1. Next, to investigate whether the amount of genetic change from standing genetic variation can help explain the extent of fitness change, we compared the number of alleles that changed their frequency during evolution with the change in growth rate (fitness) of the evolved populations. To do so, we calculated the Pearson correlation coefficient between these two metrics (paired sample correlation test). We calculated this Pearson correlation for each cut-off value for the magnitude of allele frequency change that we had used in the previous analysis of genetic change from standing variation. We did so separately for populations expanding into uniform conditions and populations expanding into a pH-gradient, as well as for sexual and asexual populations (Figure 5). Thus we calculated the correlation coefficient for four groups containing four evolved populations, combining populations with and without gene flow in a single group.

**Figure 4:**
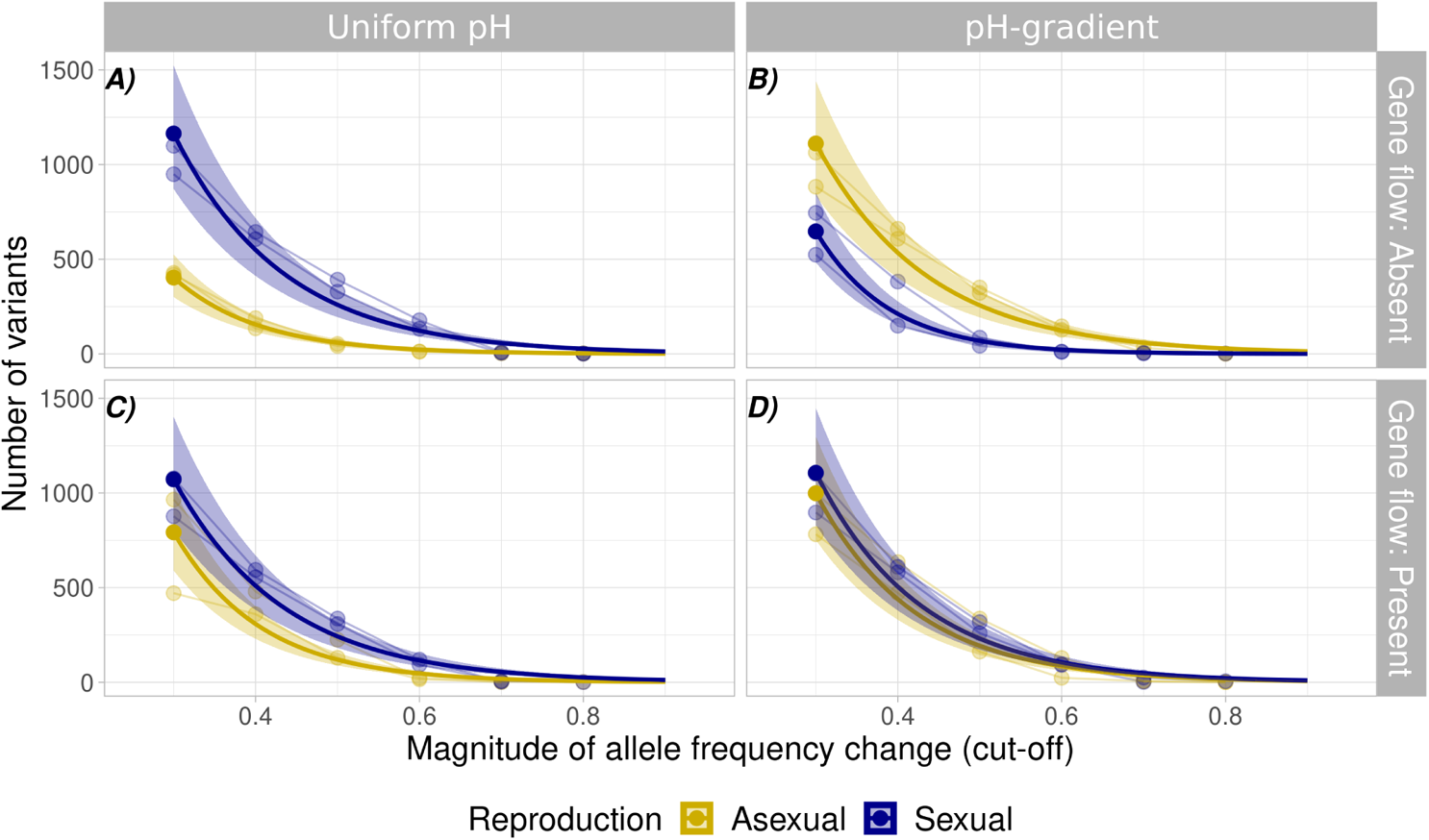
Reproduction alters genetic changes in standing genetic variation when gene flow is absent. Number of alleles that changed significantly in frequency during range expansion into a uniform environment (panel A and C) or into a pH-gradient (panel B and D). The x-axis shows the cut-off value in the magnitude of allele frequency. The y-axis shows the number of variants analyzed at each cut-off value. Circles represent allele frequency data, with thin lines connecting data from the same replicate population. Thick opaque lines show the model predictions for the best model. Shaded areas show 95%-confidence intervals from the posterior distribution. Colours represent mode of reproduction (yellow: asexual, blue:sexual).

**Figure 5:**
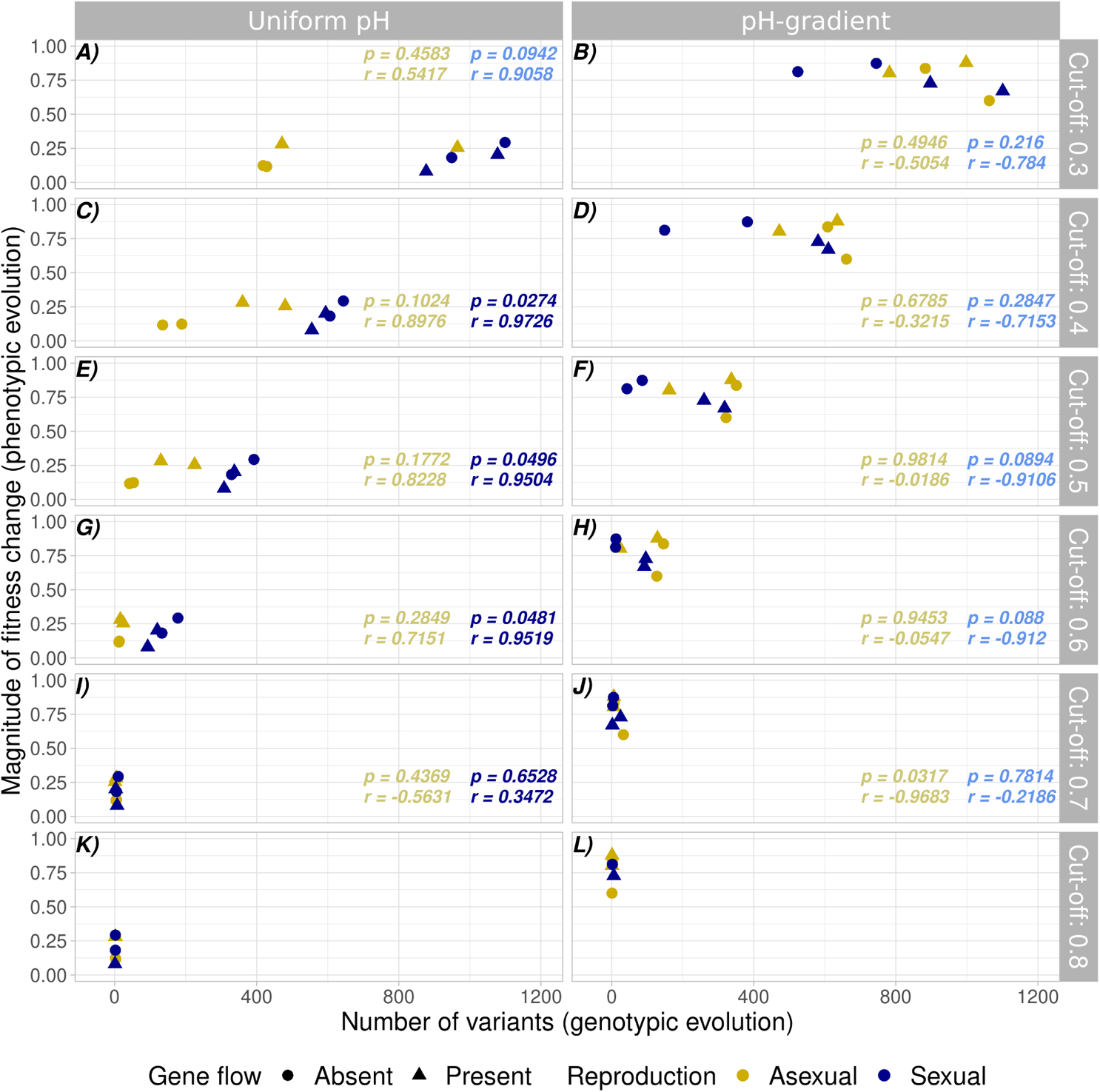
A positive correlation between fitness change and genetic change when sexual populations expand into a uniform environment. Statistical association between change in fitness and the number of allele frequency changes in standing genetic variation, for populations expanding into a uniform environment (panels A, C, E, G, I, and K) and populations expanding into a pH-gradient (panels B, D, F, H, J, and L). The x-axis shows the number of standing genetic variants that changed significantly in allele frequency. The y-axis shows the change in growth rate (fitness) of evolved populations compared to the ancestral population. Horizontal subplots show the data for different cut-off values used for the minimum change in allele frequency at the end of evolution for inclusion in the analysis. Each geometric symbol within a subplot represents a single population, with colour representing reproductive mode (yellow=asexual, blue=sexual), and shape representing gene flow (circle=gene flow absent, triangle=gene flow present). Text insets show the Pearson correlation coefficient *r* and significance p, based on a paired sample correlation test for sexual populations (blue text) and asexual populations (yellow text). Non-significant correlations are shown in light color (yellow or blue), whereas significant correlations are shown in dark color. For the most stringent cutoff value (allele frequency change >0.8), we did not calculate the correlation, because several populations did not harbour any variants that changed so dramatically in allele frequency.

For asexual populations (yellow circles and triangles), he number of alleles changing their frequency and fitness change are not correlated. In contrast, for sexual populations (blue circles and triangles), the number of alleles changing their frequency and the extent of fitness change are correlated, depending on the abiotic conditions. Specifically, for populations expanding into a uniform environment, populations in which more variants changed their frequency also increased their fitness to a greater extent (Figure 5, left panels). In contrast for populations expanding into a pH-gradient, these metrics were not correlated (Figure 5, right panels).

To explain why, in the absence of gene flow, sexual reproduction resulted in more allele frequency changes for populations expanding into a uniform environment, and fewer such changes for populations expanding into a pH-gradient, we considered the evolutionary history of our populations. Because the conditions a population experiences when expanding into a uniform environment are similar to those experienced by the ancestral population, selection on standing genetic variation may suffice for adaptation. This possibility is supported by the positive correlation between the number of genetic changes and the fitness increase for populations expanding into a uniform environment (Figure 5). Sexual recombination could help such populations adapt by bringing together adaptive alleles from different ancestor clones (Maynard-Smith, 1978). In contrast, in the pH-gradient (novel environment) new mutations or few variants from standing variation with a strong beneficial fitness effect may have been more important. In asexual populations, the lack of sexual recombination implies that any *de novo* variant or any initially rare beneficial variant from standing variation cannot be separated from its genetic background. Thus, if a beneficial variant is under strong positive selection, neutral or deleterious alleles that co-occur with it may hitch-hike to high allele frequencies. The importance of hitch-hiking is supported by the observation that in asexual populations, the number of allelic changes does not correlate with fitness (see also Figure S6), suggesting that most allelic changes don’t affect fitness strongly. In both cases, sexual reproduction may have aided adaptation, either by bringing together adaptive variants in the uniform environment, or by separating adaptive mutations from their genetic background in the pH-gradient.

When gene flow was present, we observed no clear difference in the number of variants whose frequency changed between asexual and sexual populations. The reason may be the detrimental effect of gene flow on adaptation (Lenormand, 2002). Because our experiment linked gene flow with sexual reproduction, sexual reproduction may have led to the swamping of expanding populations with maladapted alleles (Haldane and Ford, 1956; García-Ramos and Kirkpatrick, 1997; Kirkpatrick and Barton, 1997), thus counteracting the adaptive effect of selection prior to sexual reproduction. Specifically, when populations expanded their range in the presence of gene flow, the repeated bouts of sexual reproduction may have reverted the adaptive benefits of previous selection. Therefore, these populations may have undergone adaptation primarily via selection of asexual clones during periods of asexual reproduction between bouts of sexual reproduction.

### Genes involved in adaptive evolution

In our next analysis, we wanted to identify specific classes of genes that may be involved in adaptive evolution. We distinguished between genes involved in general adaptation to conditions experienced by all populations, such as the growth medium and the expanding range, and genes that are specifically involved in adaptation to our pH-gradient. For both classes of genes, we here focus on alleles that changed significantly in frequency during evolution, and report an analogous analysis of *de novo* variants in Supplementary Material section S15.

#### Changes in standing variation associated with general adaptation involve transmembrane proteins and kinase domains

Of all polymorphic genomic loci at which significant allele frequency changes occurred, 242 loci fit our general adaptation criterion. Of these 242 loci, 53 occurred in protein-coding regions, which fell into 43 different genes (Table S16). Of these 43 genes, 12 encode transmembrane proteins, four encode kinase domain proteins, three encode cyclic nucleotide-binding domain proteins, three encode zinc-fingers, and two encode cation channel proteins. All other genes either encode uncharacterized proteins (10 out of 43 genes) or fell into unique functional categories (see also Table S16).

#### Gradient-specific changes in allele frequency are associated with ion balance

We found 97,690 genomic positions for which we could quantify differences in allele frequencies through a Cochran-Mantel-Haenszel test (Methods). After Bonferroni correction, we retained 4,388 positions with significantly different allele frequencies between populations expanding into a pH-gradient and populations expanding into a uniform environment. 1,758 of these positions occurred in protein-coding regions, and fell into 636 different genes (Table S17 in the Supplementary Material). The largest subset of these genes with a functional annotation (136 genes) encode transmembrane proteins. Other highly represented groups include genes that encode kinase domain proteins (99 genes), cyclic-nucleotide binding domain proteins (39 genes), cation channel proteins (18 genes), and zinc-fingers (14 genes). The remaining genes either encode uncharacterized proteins (132 genes) or they fell into functional categories represented by only few genes. Notably, two genes, a gene encoding a cation channel family protein and a gene encoding an oxalate/formate antiporter were involved in both gradient-specific changes in allele frequency and gradient-specific *de novo* variants (Table S17 and Table S19 in the Supplementary Material).

#### General adaptation preferentially affects genes involved in DNA repair and gene expression

To find out whether genes involved in general adaptation and in gradient-specific adaptation were associated with different functions, we studied the differential enrichment of gene ontology terms in these two groups of genes, focusing on those gene ontology terms with clear differences between the groups.

Among 701 GO-terms associated with the 43 general adaptation genes, 72 terms in 13 major categories were significantly enriched after multiple testing correction (Benjamini and Hochberg, 1995; Kofler et al., 2011). Likewise, among 1928 GO-terms associated with 626 gradient-specific genes, 693 terms in 13 major categories were significantly enriched.

In Figure 6, the percentage of the 13 major categories is shown for the GO-terms associated with general adaptation (panel A) and gradient-specific adaptation (panel B). In both datasets, we found many enriched GO-terms related to metabolism, but the percentage of GO-terms did not differ significantly between the two datasets. We found only two significantly differentially enriched GO-term categories. The first comprise transcription and translation functions, which were more likely to be associated with genes involved in general adaptation (8.97% of all GO-terms) than with genes involved in gradient-specific adaptation (1.3% of all GO-terms; FDR<0.001). The second comprises mitosis, DNA repair and chromosome division, which were also more likely to be associated with genes involved in general adaptation (16.67% of all GO-terms) than in genes associated with genes gradient-specific adaptation (3.75% of all GO-terms; FDR<0.001). We thus found evidence that genes related to gene expression and genes related to cell division were under selection in all our populations, indicating that changes in such genes may be adaptive at a range edge. These observations are consistent with range expansion theory, which predicts strong selection on population growth rate at a range edge (Burton et al., 2010; Phillips et al., 2010; Shine et al., 2011).

**Figure 6:**
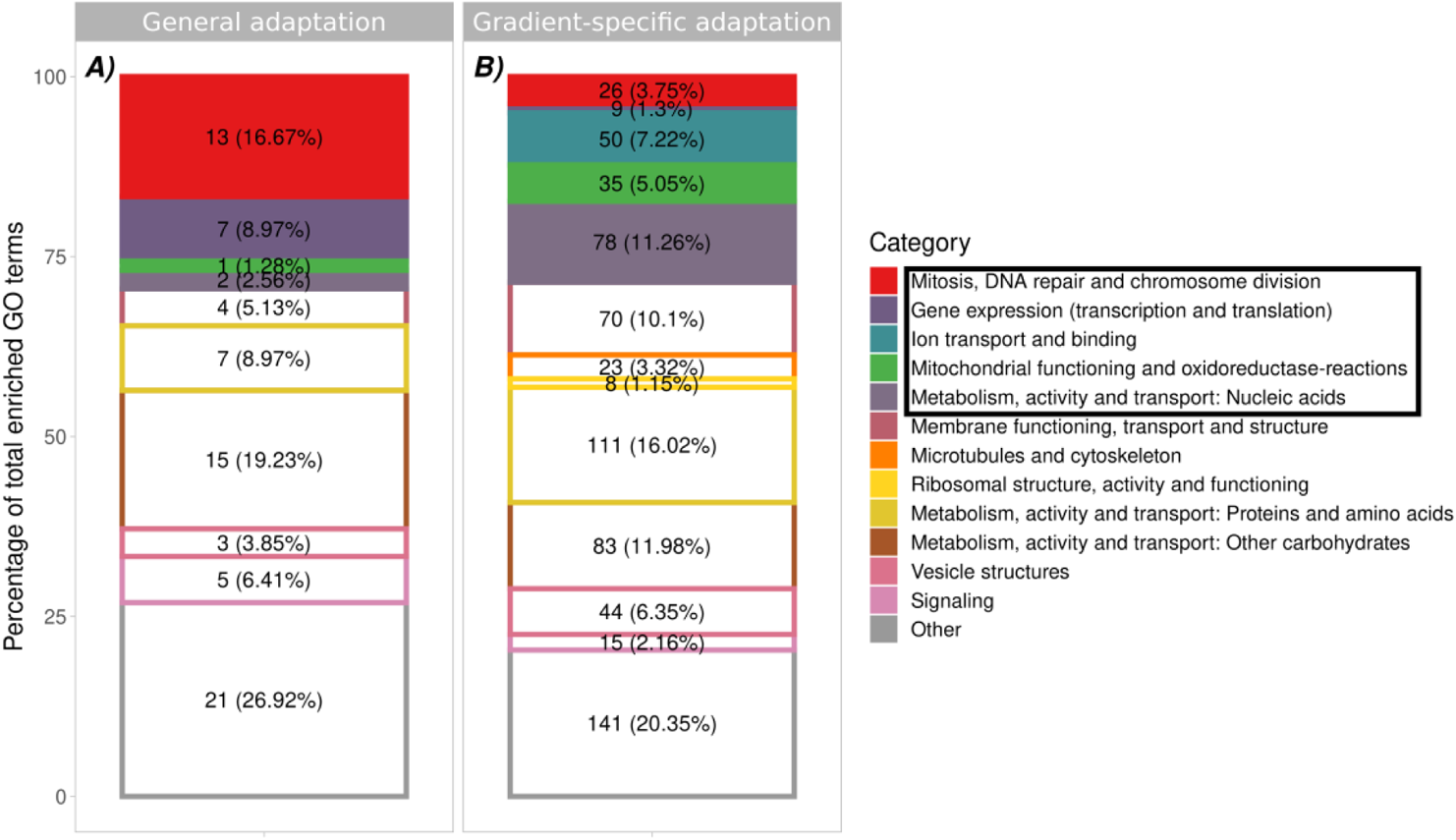
General and gradient-specific adaptations during range expansion. Partitioning of all enriched gene ontology terms (i.e. terms describing biological or molecular function of genes, as well as cellular structure) among 13 major functional categories for genes involved in general adaptations (panel A) and gradient-specific adaptations (panel B). Colours correspond to the 13 major GO categories we identified. Numbers in each rectangle represent the number and percentage of genes whose annotation fell in each category. Differentially enriched categories are displayed as filled rectangles and are boxed in the color legend. The remaining categories are displayed as open rectangles with a coloured border.

Range expansion theory also predicts strong selection for increased dispersal (Burton et al., 2010; Shine et al., 2011). Although dispersal behaviour often has a genetic basis (Saastamoinen et al., 2018), we did not observe selection on genes obviously associated with dispersal in this experiment.

Genes involved in gradient-specific adaptation were associated more often with ion binding and transport, as well as with mitochondrial functioning and oxidoreductase reactions (see figure 6). This suggests that genes coding for ion-related functions and mitochondrial-related functions may be adaptive depending on the pH environment. However, these differences were no longer significant after correcting for multiple testing. In general, genetic adaptations related to ion transport and binding may help counter osmotic stress under low pH conditions (Bremer and Krämer, 2019). Oxidoreductases play an important role in the production of reactive oxygen species (ROS), which are harmful to cells (Esterházy et al., 2008). Low pH can also contribute to elevated production of ROS (Lambert and Brand, 2004; Lemarie et al., 2011). A previous study found that selection for the capacity to metabolize ROS is an important aspect of evolution under pH-stress in phytoplankton (Lindberg and Collins, 2020). Consequently, genes associated with mitochondrial functioning and oxidoreductase reactions may help cope with the harmful consequences of ROS.

### Limitations

While our experiments identified multiple genomic changes associated with range expansions, our ability to generalize from them is subject to several limitations. The first of them is the strong selection we observed against one ancestral clone. This phenomenon required us to apply stringent criteria to identify genes under selection, and may thus have masked some signatures of general adaptation. Specifically, it limited our ability to detect polygenic adaptation of multiple weakly selected variants, and complementary but non-parallel adaptation of different genes in different populations (Stephan, 2016; Barghi et al., 2020). Future range expansion experiments should be designed with the detection of these adaptation mechanisms in mind.

A second limitation is that we could not quantify the fitness effect of individual genetic changes. The reason is that we observed many genetic changes, and that it is difficult to genetically engineer *T. thermophila* to study the effect of any one such change. A different study organism may be needed to overcome this limitation.

Thirdly, because *T. thermophila* has a large genome, we could only sequence pooled samples from a limited number of populations. One consequence is a loss of linkage information for specific variants, which limits our understanding of the role of sexual reproduction. Another consequence is limited replication of sequencing across populations, which reduces our ability to identify signatures of selection based on parallel evolution across populations. The sequencing of more populations or individuals rather than pooled populations remains an important task for future work.

## Conclusions

We showed in this experiment that sexual reproduction can alter both phenotypic evolution and the underlying genetic architecture during range expansions, depending on both the presence of a pH-gradient and the presence of long-distance gene flow. Our findings suggest that sexual reproduction may bring together adaptive variants from standing variation when populations expand in a uniform landscape. In addition, sexual reproduction may help separate adaptive from deleterious mutations and unlock the hidden benefits of rare genetic variants when populations expand their range into the novel environment of a pH-gradient. Our previous observation that clonal populations also undergo strong adaptation to a low pH environment in the absence of sexual reproduction (Moerman et al., 2020a) suggests mutations rather than standing variation may play a major role for evolutionary adaptation to the novel pH-gradient environment.

Genetic adaptation in our experiment mostly affected life-history traits and the ability to grow in the abiotic environment. Unlike previous experiments, we found no indication for the evolution of dispersal behaviour itself, neither on a genetic level or on a phenotypic level, indicating that in this experiment, the role of natural selection was more important than the role of spatial sorting.

## Acknowledgments

We thank Samuel Hürlemann, Silvana Käser and Sarah Bratschi for help with laboratory work, Vanessa Weber De Melo for aid on the nuclear separation protocol, Carla Bello and Hélène Boulain for help with the bioinformatics, as well as two anonymous reviewers for their helpful comments on a previous version of the manuscript. Funding is from the University of Zurich URPP Evolution in Action and the Swiss National Science Foundation, Grant No PP00P3 179089. This is publication ISEM-YYYY-XXX of the Institut des Sciences de l’Evolution – Montpellier. We would also like to acknowledge support by Swiss National Science Foundation grant 31003A 172887 and European Research Council Advanced Grant No. 739874.

## Data availability statement

Phenotypic data and output files from variant calling, Cochran-Mantel-Haenszel test and GO-enrichment analyses will be uploaded to Dryad upon acceptance. Raw sequence files will be uploaded to the European Sequence Archive upon acceptance. All analysis scripts and video analysis parameters will be deposited to Github on acceptance.

## Author contributions

FM, EAF, AW and FA designed the experiment. FM performed experimental work and statistical analyses. FM, FA, AW and EAF interpreted the results. FM and AW wrote the first version of the manuscript and all authors commented on and approved of the final version.

