## Supplementary Material for "Selection on growth rate and local adaptation drive genomic adaptation during experimental range expansions in the protist *Tetrahymena thermophila*"

### Supplementary Material: Experimental methodology

#### S1 Detailed experimental design

##### S1.1 Strain identities and mixing of the ancestral population

We used in this experiment four ancestral clones, which we obtained from the Tetrahymena Stock Center (`https://tetrahymena.vet.cornell.edu/`). These ancestor clones were strain B2086.2 (Research Resource Identifier TSC_SD00709), strain CU427.4 (TSC_SD00715), strain CU428.2 (TSC_SD00178) and strain SB3539 (TSC_SD00660). We created the ancestral population by mixing these four ancestor clones together. To do so, we first grew the four ancestor clones separately to equilibrium density and measured cell counts for each of the four ancestors. Subsequently, we mixed equal volumes of the resulting four cultures. Because each clone may grow to a different equilibrium density, the clones may not be represented by equal cell numbers in the ancestral population. The starting proportions of the four clones in the ancestral population (i.e. the proportion of cells of the ancestral population originating from each of the four clones) are shown in Table S1. We then used this ancestral population to initiate the range expansion experiment described in the main text (section Experimental evolution).

| Clone | Starting proportion |
|---|---|
| **Clone B2086.2** | 0.3402 |
| **Clone CU427.4** | 0.1287 |
| **Clone CU428.2** | 0.2595 |
| **Clone SB3539** | 0.2716 |

Table S1 Starting proportions of cells in our ancestral population that originated from each of the indicated ancestral clones.

##### S1.2 Two-patch landscapes

The two-patch landscapes we used consist of two 25 mL Sarstedt tubes, each filled with 15 mL of modified Neff medium (Cassidy-Hanley, 2012). The Sarstedt tubes are connected by an 8 cm long silicone tube (inner diameter 4 mm), that can be opened or closed using a plastic clamp, in order to allow cells to disperse between the two Sarstedt tubes.

##### S1.3 Detailed explanation of experimental handling

During the range expansion experiment, we repeated a cyclic handling procedure every 14 days. This procedure consisted of three dispersal events (days 1, 3, and 5 of the 14 day cycle), followed by a gene flow and sexual reproduction event in the relevant treatments (day 8 of the 14 day cycle), and finally two additional dispersal events (days 10, and 12 of the 14 day cycle). We here described in more detail how we performed the dispersal, gene flow and sexual reproduction events during experimental evolution.

A dispersal event consisted of opening the plastic clamps of the two-patch landscapes for one hour, which allowed cells to swim from the home patch to the target patch. After dispersal, we measured cell densities in the home and target patches using a Leica M165FC stereomicroscope with a top-mounted Hamamatsu Orca Flash 4.0 camera andusing an established video analysis method (Pennekamp et al., 2015) and a Leica M165FC stereomicroscope with top-mounted Hamamatsu Orca Flash 4.0 camera. We then prepared 40 new two-patch landscapes.

For all populations that had successfully dispersed into the target patch, we transferred the content of the target patch to the home patch of the new two-patch landscape. For all other populations, we transferred the content of the home patch to the new two-patch landscape. For populations designated for range expansion in a pH-gradient, we gradually lowered the pH of the Neff medium in the new two-patch landscapes. Specifically, we lowered the pH from 6.5 in steps of 0.5 after every 2-3 successful dispersal events, until a minimal value of pH 4.0 had been reached. From that time onward, we kept the populations at pH 4.0 for the remainder of the range expansion experiment. Although genetic drift may play an important role during range expansions (Hallatschek et al., 2007; Excoffier et al., 2009), it is not likely to play a major role on the time scale of in our experiments. Firstly, population densities in our experiments were high ($10^4 - 10^6$ individuals). Secondly, between 1% and 20% of our populations typically dispersed, such that dispersal never led to extreme population bottlenecks . Therefore, genetic drift is negligible on the timescale of this experiment (Hartl and Clark, 2006).

To implement long-distance gene flow from the range core to the range edge, we replaced 1.5 mL of culture in the appropriate populations with 1.5 mL of a replicate of the ancestral population. We created this replicate by mixing the four ancestral clones in the same proportions as we had used to create the ancestral population from which we had started the experiment.

Following a gene flow event, we induced sex in populations designated for sexual reproduction. *T. thermophila* only mates when starved (Lynn and Doerder, 2012). Therefore, we transferred all populations after a gene flow event to starvation medium, and incubated them on a shaker rotating at 120 rpm for 36 hours. Subsequently, we removed populations designated for sexual reproduction from the shaker, but kept populations designated for asexual reproduction on the shaker, because the shaking prevents cells from mating. We then allowed cells to mate overnight, before transferring all populations back from starvation medium to modified Neff medium.

We repeated these described handling steps during every 14 day cycle, with two exceptions. In the first cycle, we did not perform a dispersal event on day 1, because the experimental populations had only just been inoculated, and therefore population densities were too low for measurable dispersal to occur. Secondly, during the fifth and last 14 day cycle, we did not initiate an additional gene flow and sexual reproduction event.

##### S1.4 Visual representation of experimental evolution phase

##### S1.5 Population Growth assessments

After experimental evolution and common garden treatment (see Materials and Methods in the main text), we performed population growth assays to compare how population growth rate differed between the ancestral population and the evolved populations. To do so, we prepared Sarstedt tubes containing modified Neff medium with the pH adjusted to eight different values (pH 6.5, 6.0, 5.5, 5.0, 4.5, 4.0, 3.5 and 3.0) for the ancestral population and for each evolved populations. We then inoculated the Sarstedt tubes either with 100 µL of culture from the ancestral population or from one of the evolved populations. We grew these populations for 12 days, during which we sampled populations twice on day one and two as well as once on every subsequent day. We then applied existing video analysis methods (Pennekamp et al., 2015), using a Leica M165FC stereomicroscope with top-mounted Hamamatsu Orca Flash 4.0 camera to assess population densities.

After video analysis, we analyzed population growth using a continuous-time version of the Beverton-Holt population growth model (Beverton and Holt, 1993). This model is well suited for microcosm data and has biologically interpretable parameters (Thieme, 2003; Fronhofer

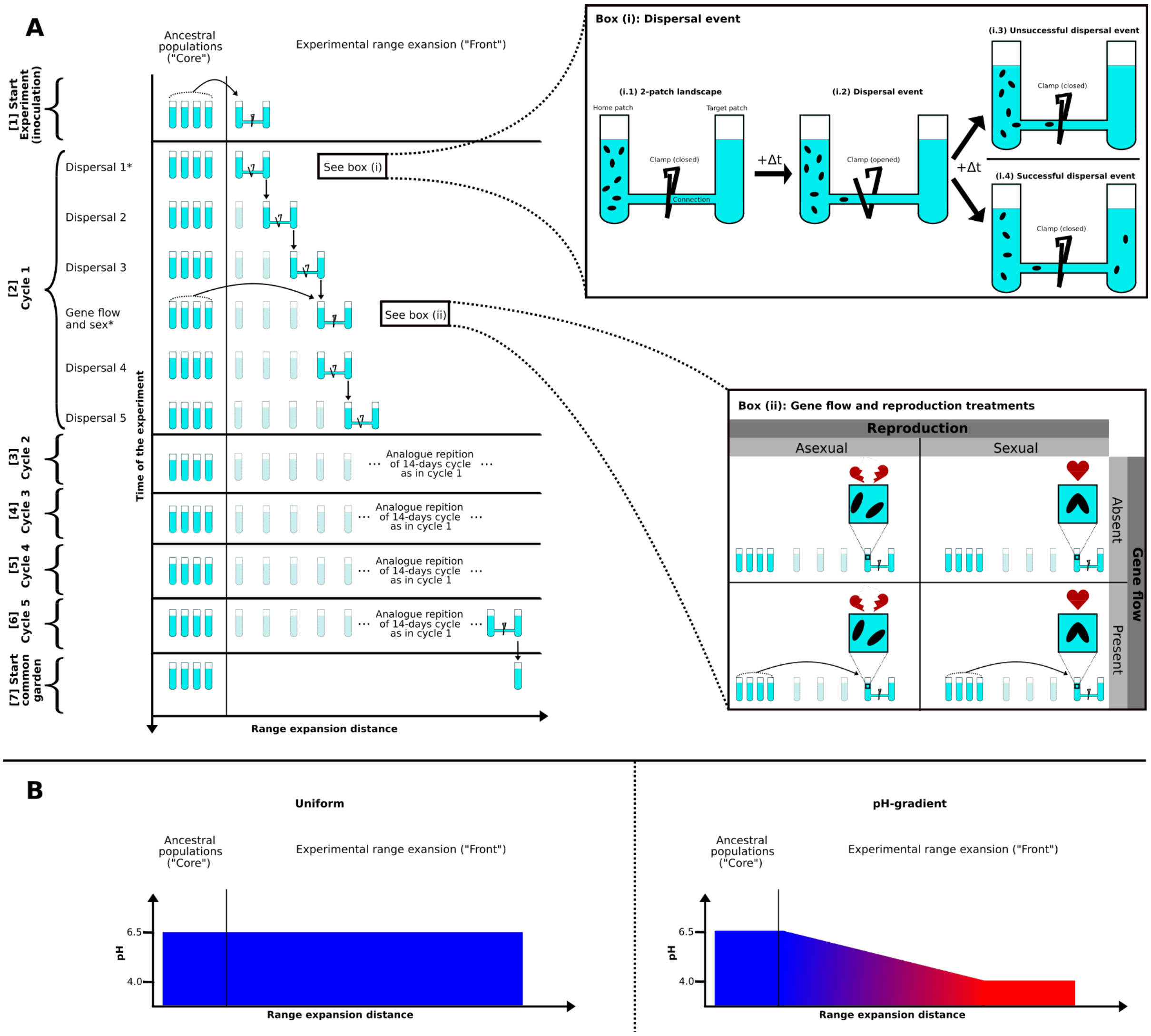


Figure S1: **Schematic representation of the experimental range expansion.** Adapted from Moerman et al. (2020b). Panel A shows the timeline of the experiment, with the y-axis depicting time, and the x-axis the distance by which populations expanded during range expansion. Box (i) shows in more detail the experimental approach involved in a dispersal step, and box (ii) the treatments related to gene flow and reproduction. Panel B shows the abiotic conditions during range expansion into a pH-gradient and a uniform environment.

et al., 2018). The Beverton-Holt model is given by the equation

$$\frac{dN}{dt} = \left(\frac{r_0 + d}{1 + \alpha N} - d\right) N, \tag{2}$$

where the intraspecific competitive ability ($\alpha$) is given by

$$\alpha = \frac{r_0}{\widehat{N} d}. \tag{3}$$

In these equations, $r_0$ is the intrinsic population growth rate, $N$ is the population size, $\alpha$ is the intraspecific competitive ability, $\widehat{N}$ is the equilibrium population density and $d$ is the death rate of the population. We used a Bayesian approach adapted from Rosenbaum et al. (2019) for parameter estimation. Model code is available at `https://zenodo.org/record/2658131`.

### S2 Nuclear separation

We here describe the detailed protocol used to separate macronuclei and micronuclei of our evolved populations. We first list the three media used in the protocol with all required reagents. We then describe in detail how we prepared and stored the media, and how we performed nuclear separation for our populations.

#### S2.1 Media reagents for nuclear separation

##### Reagents for medium A

| Reagent | Concentration | Amount/1 L |
|---|---:|---:|
| Sucrose | 0.1 M | 34.23 g |
| $MgCl_2(*H_2O_6)$ | 2 mM | 0.407 g |
| Gum arabic | 4% | 40 g |
| Tris pH 7.5 (Tris base, with pH adjusted) | 10 mM | 1.211 g |
| Iodoacetamide | 1mM | 0.185 g |
| Butyric acid | 10 mM | 0.881 g / 0.918 µL |

Table S2 Reagents required to prepare of 1 L of medium A.

##### Reagents for nuclei was buffer

| Reagent | Concentration | Amount/1 L |
|---|---:|---:|
| Sucrose | 0.25 M | 85.574 g |
| Tris pH 7.5 (Tris base, with pH adjusted) | 10 mM | 1.211 g |
| $CaCl_2$ | 3 mM | 0.441 g |
| $MgCl_2$ | 1 mM | 0.2033 g |
| Iodoacetamide | 1mM | 0.185 g |
| Butyric acid | 10 mM | 0.881 g / 0.918 µL |

Table S3 Reagents required to prepare 1 L of nuclei wash buffer.

##### Ingredients of SyBr-Green Tris-HCl

| Reagent | Amount |
|---|---:|
| 10 mM Tris-HCl (pH 7.5) | 50 mL |
| SyBr-Green 10.000X concentrated | 50 µL |

Table S4 Ingredients required to prepare the SyBr-Green Tris-HCl solution for nuclear staining.

#### S2.2 Nuclear separation protocol

Prior to nuclear separation, we prepared the required media (medium A and nuclei wash buffer). Briefly, to prepare medium A, we mixed 34.23 g of sucrose, 0.407 g of $MgCl_2(\times H_2O_6)$, 40 g gum arabic, and 0.185 g iodoacetamide in a 1 liter Schott bottle containing 800 mL of ultra-pure water (i.e. water filtered using a Barnstead Waterpure filtration system to remove organic and inorganic compounds; see `https://assets.thermofisher.com/TFS-Assets/LED/manuals/D01316~.pdf` for the manufacturer protocol). We then added 0.918 µL butyric acid and 1.211 g of Tris at pH 7.5. We prepared this pH-adjusted Tris by first creating a stock solution from Tris-Base, and adjusted the pH to 7.5 using 1M HCl. After thoroughly mixing all ingredients of medium A, we then filled the Schott bottle with ultrapure water, until the total volume was 1 L. After mixing again, we then transferred 40 mL of medium A to 50 mL Falcon tubes, and stored the Falcon tubes in a −20 °C freezer until needed. One Falcon tube suffices for the nuclear separation of one population.

We prepared the nuclei wash buffer by mixing 85.574 g of sucrose, 0.441 g $CaCl_2$, 0.2033 g $MgCl_2$ and 0.185 g iodoacetamide in a 1 L Schott bottle containing 800 mL of ultrapure water. We then added 0.918 µL butyric acid and 1.211 g of Tris at pH 7.5. We then filled the bottle with ultrapure water until the total volume was 1 L, and mixed thoroughly. After mixing, we transferred 15 mL of nuclei wash buffer to 15 mL Falcon tubes, and stored these tubes in a −20 °C freezer until needed.

Before starting the nuclear separation protocol, we moved the required number of Falcon tubes to a 4 °C refrigerator to thaw overnight. We then prepared medium B by adding 50 µL of octanol to the Falcon tubes containing medium A. After adding the octanol, we thoroughly mixed the Falcon tubes by vortexing, and stored them on ice.

Next, we determined the population density of all populations from which we wanted to extract DNA. To do so, we used an existing video analysis method (Pennekamp et al., 2015) to measure the number of cells in a small volume of culture for each population, using a Leica M165FC stereomicroscope with top-mounted Hamamatsu Orca Flash 4.0 camera. One Falcon tube of medium B suffices for nuclear separation of approximately $4 \times 10^7$ cells of *T. thermophila*. Therefore, we collected for every population the volume of culture containing a total of $4 \times 10^7$ cells, and transferred this volume to 50 mL Falcon tubes. We then centrifuged these Falcon tubes at 2000 g and 4 °C for 5 minutes, using a Sigma 3-16PK centrifuge. After centrifugation, we decanted the supernatant, and resuspended the cells in 5 mL of medium B. We then transferred the suspended cells to a glass homogenizer dounce (Kimble 15 mL tissue grinder; product ID: 885300-0015). Subsequently, we ruptured cells and separated their nuclei using 40 pushes of the glass dounce. After cell disruption, we verified that cells had been disrupted and that nuclei were separated in at least 90% of the cells (i.e. the smaller micronucleus was dislodged from the ”cup” next to the larger macronucleus). To do so, we stained a small droplet of ruptured cells using SyBr-green dye to colour the DNA. We then visually inspected the droplet under a Leica M165FC stereomicroscope with a green fluorescence filter to verify that macronuclei and micronuclei were separated.

After cell disruption, we transferred the cell material to the remaining 35 mL of medium B in the Falcon tube, and thoroughly mixed the content of the Falcon tubes. We then centrifuged the Falcon tubes (2000 g at 4 °C for 5 minutes) using a Sigma 3-16PK centrifuge to pellet the macronuclei. We decanted the supernatant (containing micronuclei, unpelleted macronuclei, and other cell material) in a new 50 mL Falcon tube, and resuspended the pellet in 1 mL of nuclei wash buffer (pellet P1). We then transferred the resuspended pellet to a 1.5 mL Eppendorf tube, and immediately stored the Eppendorf tube on ice. Next, we mixed the supernatant by vortexing, and centrifuged it again at a higher speed (2300 g at 4 °C for 5 minutes; Sigma 3-16PK centrifuge) to pellet the remaining macronuclei. We then again decanted the supernatant in a new 50 mL Falcon tube, and resuspended the second pellet (pellet P2) in 1 mL of nuclei wash buffer. We transferred the resuspended P2 pellet to a 1 mL Eppendorf tube, which we then stored on ice. Subsequently, we again mixed the remaining supernatant by vortexing, and centrifuged it at increased speed (2500 g at 4 °C for 5 minutes; Sigma 3-16PK centrifuge). We then decanted the supernatant in a waste disposal collection bottle, and resuspended the pellet in 1 mL of nuclei wash buffer (pellet P3). Subsequently, we transferred the resuspended P3 pellet to a 1 mL Eppendorf tube which we stored on ice. After nuclear separation, we immediately started the DNA extraction from the resuspended pellets containing the macronuclei.

### Supplementary Material: Bioinformatic methodology

#### S3 DNA quality assessment

We verified the quality and quantity of our DNA samples after DNA extraction using a Nanodrop spectrophotometer (model ND-1000). Specifically, we aimed to verify that the 260 nm/280 nm and 260 nm/230 nm ratios were within acceptable values, and ranked the pellets by DNA quality and quantity. We used the following criteria to rank the quality of the DNA samples. We chose for sequencing the DNA extracted from the pellet with a 260 nm/280 nm ratio closest to 1.8, a 260 nm/230 nm ratio larger than 2, and the highest concentration of DNA among all three pellets. If these criteria were similar between the macronuclear pellets, we always chose the DNA extracted from the first pellet (P1).

#### S4 Read mapping and genome coverage statistics

We here list for all our sequenced populations the relevant statistics related to read mapping against the reference genome, and the genome coverage we obtained.

#### S4.1 Read mapping statistics

| Population | History | Reads | Mapped reads | Percentage mapped | Abiotic conditions | Gene flow | Reproduction |
|---|---|---|---|---|---|---|---|
| B2086.2 | Ancestral | 88538723 | 87177869 | 98.46 | Ancestral | Ancestral | Ancestral |
| CU427.4 | Ancestral | 60231352 | 58955342 | 97.88 | Ancestral | Ancestral | Ancestral |
| CU428.2 | Ancestral | 121454032 | 119887118 | 98.71 | Ancestral | Ancestral | Ancestral |
| SB3539 | Ancestral | 55923710 | 54488844 | 97.43 | Ancestral | Ancestral | Ancestral |
| EVO4 | Evolved | 151091447 | 148015322 | 97.96 | pH-gradient | Present | Sexual |
| EVO5 | Evolved | 61936120 | 60664281 | 97.95 | pH-gradient | Present | Sexual |
| EVO8 | Evolved | 66688733 | 65342004 | 97.98 | pH-gradient | Present | Asexual |
| EVO9 | Evolved | 74980890 | 73779386 | 98.4 | pH-gradient | Present | Asexual |
| EVO11 | Evolved | 81714042 | 79260522 | 97.00 | pH-gradient | Absent | Sexual |
| EVO14 | Evolved | 145705318 | 142906377 | 98.08 | pH-gradient | Absent | Sexual |
| EVO18 | Evolved | 58985398 | 57659400 | 97.75 | pH-gradient | Absent | Asexual |
| EVO20 | Evolved | 65135354 | 63837265 | 98.01 | pH-gradient | Absent | Asexual |
| EVO23 | Evolved | 120307664 | 116904548 | 97.17 | Uniform | Present | Sexual |
| EVO24 | Evolved | 67309446 | 65539280 | 97.37 | Uniform | Present | Sexual |
| EVO29 | Evolved | 113384019 | 111097232 | 97.98 | Uniform | Present | Asexual |
| EVO30 | Evolved | 50504039 | 49125464 | 97.27 | Uniform | Present | Asexual |
| EVO33 | Evolved | 104327400 | 101909411 | 97.68 | Uniform | Absent | Sexual |
| EVO35 | Evolved | 73190459 | 71198417 | 97.28 | Uniform | Absent | Sexual |
| EVO36 | Evolved | 62024246 | 60916336 | 98.21 | Uniform | Absent | Asexual |
| EVO39 | Evolved | 78497531 | 75953004 | 96.76 | Uniform | Absent | Asexual |

Table S5 Read mapping statistics following mapping of the 2×150 bp paired-end Illumina reads. For each of the ancestor clones and evolved populations, we list the total number of reads (Reads), the number of reads that mapped to the reference genome (Mapped reads), and the percentage of total reads that mapped to the reference genome (Percentage mapped reads).

#### S4.2 Coverage statistics

| Population | History | Mean coverage | St. Dev of coverage | Abiotic conditions | Gene flow | Reproduction |
|---|---|---:|---:|---|---|---|
| B2086.2 | Ancestral | 118.755 | 43.597 | Ancestral | Ancestral | Ancestral |
| CU427.4 | Ancestral | 81.246 | 38.903 | Ancestral | Ancestral | Ancestral |
| CU428.2 | Ancestral | 161.785 | 48.709 | Ancestral | Ancestral | Ancestral |
| SB3539 | Ancestral | 199.782 | 53.243 | Ancestral | Ancestral | Ancestral |
| EVO4 | Evolved | 82.371 | 39.575 | pH-gradient | Present | Sexual |
| EVO5 | Evolved | 88.575 | 40.435 | pH-gradient | Present | Sexual |
| EVO8 | Evolved | 100.868 | 40.768 | pH-gradient | Present | Asexual |
| EVO9 | Evolved | 107.539 | 47.846 | pH-gradient | Present | Asexual |
| EVO11 | Evolved | 193.385 | 56.618 | pH-gradient | Absent | Sexual |
| EVO14 | Evolved | 78.104 | 28.078 | pH-gradient | Absent | Sexual |
| EVO18 | Evolved | 86.787 | 41.000 | pH-gradient | Absent | Asexual |
| EVO20 | Evolved | 158.747 | 55.337 | pH-gradient | Absent | Asexual |
| EVO23 | Evolved | 89.327 | 43.839 | Uniform | Present | Sexual |
| EVO24 | Evolved | 150.726 | 46.979 | Uniform | Present | Sexual |
| EVO29 | Evolved | 66.802 | 39.701 | Uniform | Present | Asexual |
| EVO30 | Evolved | 138.968 | 45.759 | Uniform | Present | Asexual |
| EVO33 | Evolved | 94.284 | 41.506 | Uniform | Absent | Sexual |
| EVO35 | Evolved | 83.033 | 37.243 | Uniform | Absent | Sexual |
| EVO36 | Evolved | 103.031 | 42.444 | Uniform | Absent | Asexual |
| EVO39 | Evolved | 73.021 | 36.142 | Uniform | Absent | Asexual |

Table S6 Genome coverage statistics after sequencing the populations using 2×150 bp paired end Illumina reads. For each of the ancestor clones and evolved populations, we list the mean coverage across the entire genome (Mean coverage) and the standard deviation in coverage across the entire genome (St. Dev of coverage).

### S5 Calculation of expected allele frequencies in ancestral population

Before we could assess whether there was a change in allele frequency for any one evolved population, we needed to calculate the expected allele frequency in the ancestral population. Cells of *T. thermophila* have a highly polyploid macronucleus (n=45), and during asexual divisions the macronuclear chromosomes divide randomly (Lynn and Doerder, 2012; Ruehle et al., 2016). This means that a locus for which a clone is heterozygous in the micronucleus can be present in zero to 45 chromosome copies of the macronuclear DNA of a single cell. We therefore first needed to calculate, for each of the four ancestral clones, the allele frequency at all genome positions in which we observed a difference from the reference genome. Subsequently, we calculated a weighted mean of these four allele frequencies, using the starting proportion of the four ancestral clones in the ancestral population (Table S1). These weighted allele frequencies thus represent the expected allele frequencies of all variants that were initially present in the ancestral population.

### S6 Quality filtering of genetic variants

We here describe the criteria we used to filter candidate genomic sites (positions) for genetic changes in standing variation and for *de novo* variants. Specifically, we aimed to exclude candidate sites with poor sequence quality, low sequence coverage, or sequencing artefacts.

#### S6.1 Quality filtering of candidate sites for genetic changes from standing variation

We had identified candidate sites for genetic change from standing variation as those sites where genetic variation was present in the ancestral population. We then performed a quality filtering for these candidate sites based on the following criteria. Firstly, we removed any sites where at least one of the 20 populations had an extremely low ($<$30) number of high quality reads, thus excluding genomic regions with very low sequence coverage. Secondly, we removed sites with an extremely high ($>$300) number of high quality reads, which could be caused by sequencing artefacts, such as duplicated segments in the sequenced populations that are not present in the reference genome and may falsely map to the same genomic region. Thirdly, to reduce the incidence of false positive variant alleles, we removed any positions where only a single read differed from the reference genome in all 20 populations. Lastly, we considered only genomic positions for which we had high quality data (QUAL score $\leq$ 30) for all four ancestral clones, as well as for the 16 evolved populations.

#### S6.2 Quality filtering of candidate sites for genetic changes through *de novo* variants

We had identified candidate sites for genetic change through *de novo* variants as those sites where a genetic variant that was not present in any of the four ancestor clones occurred in at least one evolved population. From the resulting collection of such variants, we removed variants with few ($<$30) number of high quality reads, as well as variants with a low quality score (QUAL $<$30). We then filtered the remaining variants by removing variants where only a single read differed from the ancestral clones, in order to exclude false positive *de novo* variants caused by sequencing errors.

### S7 Clonal selection of clone SB3539

#### S7.1 Derived alleles in sequenced populations

| Population | History | SNPs | INDELs | Transitions | Transversions | Transition/Transversion ratio | Abiotic conditions | Gene flow | Reproduction |
|---|---|---|---|---|---|---|---|---|---|
| B2086.2 | Ancestral | 19140 | 9328 | 8204 | 10981 | 0.7471 | Ancestral | Ancestral | Ancestral |
| CU427.4 | Ancestral | 23201 | 7848 | 10088 | 13169 | 0.766 | Ancestral | Ancestral | Ancestral |
| CU428.2 | Ancestral | 12983 | 9677 | 5293 | 7737 | 0.6841 | Ancestral | Ancestral | Ancestral |
| SB3539 | Ancestral | 640765 | 80210 | 310239 | 332015 | 0.9344 | Ancestral | Ancestral | Ancestral |
| EVO4 | Evolved | 8510 | 8302 | 3394 | 5154 | 0.659 | pH-gradient | Present | Sexual |
| EVO5 | Evolved | 23494 | 8944 | 10198 | 13349 | 0.764 | pH-gradient | Present | Sexual |
| EVO8 | Evolved | 23306 | 10906 | 10057 | 13293 | 0.757 | pH-gradient | Present | Asexual |
| EVO9 | Evolved | 20477 | 9127 | 8829 | 11689 | 0.755 | pH-gradient | Present | Asexual |
| EVO11 | Evolved | 23168 | 11609 | 9869 | 13354 | 0.739 | pH-gradient | Absent | Sexual |
| EVO14 | Evolved | 12011 | 12020 | 4769 | 7288 | 0.654 | pH-gradient | Absent | Sexual |
| EVO18 | Evolved | 24468 | 8811 | 10620 | 13896 | 0.764 | pH-gradient | Absent | Asexual |
| EVO20 | Evolved | 23600 | 8632 | 10721 | 13384 | 0.801 | pH-gradient | Absent | Asexual |
| EVO23 | Evolved | 13277 | 9534 | 5455 | 7870 | 0.693 | Uniform | Present | Sexual |
| EVO24 | Evolved | 23136 | 9231 | 10036 | 13146 | 0.763 | Uniform | Present | Sexual |
| EVO29 | Evolved | 13264 | 10840 | 5447 | 7860 | 0.693 | Uniform | Present | Asexual |
| EVO30 | Evolved | 24581 | 9730 | 10592 | 14041 | 0.754 | Uniform | Present | Asexual |
| EVO33 | Evolved | 14776 | 9452 | 6161 | 8667 | 0.711 | Uniform | Absent | Sexual |

| | | | | | | | | | |
|---|---|---|---|---|---|---|---|---|---|
| EVO35 | Evolved | 22484 | 9649 | 9777 | 12745 | 0.767 | Uniform | Absent | Sexual |
| EVO36 | Evolved | 25608 | 11874 | 10911 | 14745 | 0.740 | Uniform | Absent | Asexual |
| EVO39 | Evolved | 25158 | 12406 | 10727 | 14492 | 0.740 | Uniform | Absent | Asexual |

Table S7 Derived variants found after variant calling. For each of the ancestor clones and evolved populations, we list the number of SNPs that deviate from the reference genome (SNPs), the number of indels that deviate from the reference genome (INDELs), the number of transitions found among all derived SNPs (Transitions), the number of transversion found among all derived SNPS (Transversion) and the transition to transversion ratio. Note the high number of derived SNPs and indels for clone SB3539, as well as the noticeably higher transition to transversion ratio for this clone.

### S8 Cochran-Mantel-Haenszel test

#### S8.1 Pairing of populations

| Comparison number | Population 1 (Gradient) | Population 2 (Uniform) | Gene flow | Reproduction |
|---:|---|---|---|---|
| 1 | EVO4 | EVO23 | Present | Sexual |
| 2 | EVO5 | EVO24 | Present | Sexual |
| 3 | EVO8 | EVO29 | Present | Asexual |
| 4 | EVO9 | EVO30 | Present | Asexual |
| 5 | EVO11 | EVO33 | Absent | Sexual |
| 6 | EVO14 | EVO35 | Absent | Sexual |
| 7 | EVO18 | EVO36 | Absent | Asexual |
| 8 | EVO19 | EVO39 | Absent | Asexual |

Table S8 All population comparisons used in the Cochran-Mantel-Haenszel test to identify differential selection in populations expanding into a uniform environment and populations expanding into a gradient.

### Supplementary Material: Phenotypic data and results

### S9 Intrinsic growth rate of evolved populations

| ID | Abiotic conditions | Reproduction | Gene flow | $r_0$ | $r_0$ change | Included in genetic analysis |
|---|---|---|---|---|---|---|
| EVO1 | pH-gradient | Sexual | Present | 0.0402 | 0.0627 | no |
| EVO2 | pH-gradient | Sexual | Present | 0.0603 | 0.6496 | no |
| EVO4 | pH-gradient | Sexual | Present | 0.0611 | 0.6700 | yes |
| EVO5 | pH-gradient | Sexual | Present | 0.0637 | 0.7276 | yes |
| EVO6 | pH-gradient | Asexual | Present | 0.0686 | 0.8355 | no |
| EVO7 | pH-gradient | Asexual | Present | 0.0669 | 0.7990 | no |
| EVO8 | pH-gradient | Asexual | Present | 0.0671 | 0.8029 | yes |
| EVO9 | pH-gradient | Asexual | Present | 0.0707 | 0.8783 | yes |
| EVO10 | pH-gradient | Asexual | Present | 0.0658 | 0.7747 | no |
| EVO11 | pH-gradient | Sexual | Absent | 0.0705 | 0.8735 | yes |
| EVO13 | pH-gradient | Sexual | Absent | 0.0670 | 0.8019 | no |
| EVO14 | pH-gradient | Sexual | Absent | 0.0675 | 0.8120 | yes |
| EVO18 | pH-gradient | Asexual | Absent | 0.0687 | 0.8366 | yes |
| EVO20 | pH-gradient | Asexual | Absent | 0.0583 | 0.6000 | yes |
| EVO21 | Uniform | Sexual | Present | 0.1033 | 0.0124 | no |

| | | | | | | |
|---|---|---|---|---|---|---|
| EVO22 | Uniform | Sexual | Present | 0.1077 | 0.0728 | no |
| EVO23 | Uniform | Sexual | Present | 0.1178 | 0.2030 | yes |
| EVO24 | Uniform | Sexual | Present | 0.1082 | 0.0802 | yes |
| EVO25 | Uniform | Sexual | Present | 0.1079 | 0.0761 | no |
| EVO26 | Uniform | Asexual | Present | 0.1206 | 0.2362 | no |
| EVO27 | Uniform | Asexual | Present | 0.1086 | 0.0856 | no |
| EVO28 | Uniform | Asexual | Present | 0.1116 | 0.1244 | no |
| EVO29 | Uniform | Asexual | Present | 0.1222 | 0.2551 | yes |
| EVO30 | Uniform | Asexual | Present | 0.1245 | 0.2823 | yes |
| EVO31 | Uniform | Sexual | Absent | 0.1103 | 0.1075 | no |
| EVO32 | Uniform | Sexual | Absent | 0.1073 | 0.0672 | no |
| EVO33 | Uniform | Sexual | Absent | 0.1254 | 0.2930 | yes |
| EVO34 | Uniform | Sexual | Absent | 0.1131 | 0.1443 | no |
| EVO35 | Uniform | Sexual | Absent | 0.1161 | 0.1819 | yes |
| EVO36 | Uniform | Asexual | Absent | 0.1115 | 0.1231 | yes |
| EVO37 | Uniform | Asexual | Absent | 0.1029 | 0.0079 | no |
| EVO38 | Uniform | Asexual | Absent | 0.0782 | -0.3881 | no |
| EVO39 | Uniform | Asexual | Absent | 0.1110 | 0.1162 | yes |
| EVO40 | Uniform | Asexual | Absent | 0.1089 | 0.0888 | no |

Table S9 Phenotypic change in evolved populations. For each of the evolved populations, we list the intrinsic growth rate ($r_0$), as well as the change in $r_0$ compared to the ancestral population ($r_o$ change), calculated as the logarithm (base 2) of the ratio between the intrinsic growth rate of the evolved population and the intrinsic growth rate of the ancestral population. The column ”Included in genetic analysis” indicates whether the population was selected for sequencing or not.

### S10 Correlation between intrinsic growth rate ($r_0$) and movement metrics.

We chose populations for genetic analysis based on their change in intrinsic growth rate ($r_0$). This procedure could have created a bias in our results, if the intrinsic growth rate shows a trade-off with another trait, most notably movement. To determine if such a trade-off exists, we calculated the Pearson correlation coefficient (paired sample correlation test) between the intrinsic growth rate $r_0$ on the one hand, and two movement metrics (gross swimming speed and turning speed) on the other hand. More specifically, we calculated this coefficient at three specific time points during a population's growth, i.e., 1) at the first time point of the measured growth time series (i.e., at low population density), 2) at the inflection point (i.e. at the point where the population density was closest to 50 % of the equilibrium density; medium density), and 3) at the point where the population first approached its equilibrium density (i.e. at $>$ 99 % of the equilibrium density; high density). To account for potential phenotypic plasticity caused by variation in the pH of the environment, we calculated the correlation separately for populations growing in medium with pH 6.5, and for populations growing in medium with pH 4.0. The intrinsic rate of increase ($r_0$) was not significantly correlated with either the gross movement speed (figure S2) or with the turning speed (figure S3) in any of our tests.

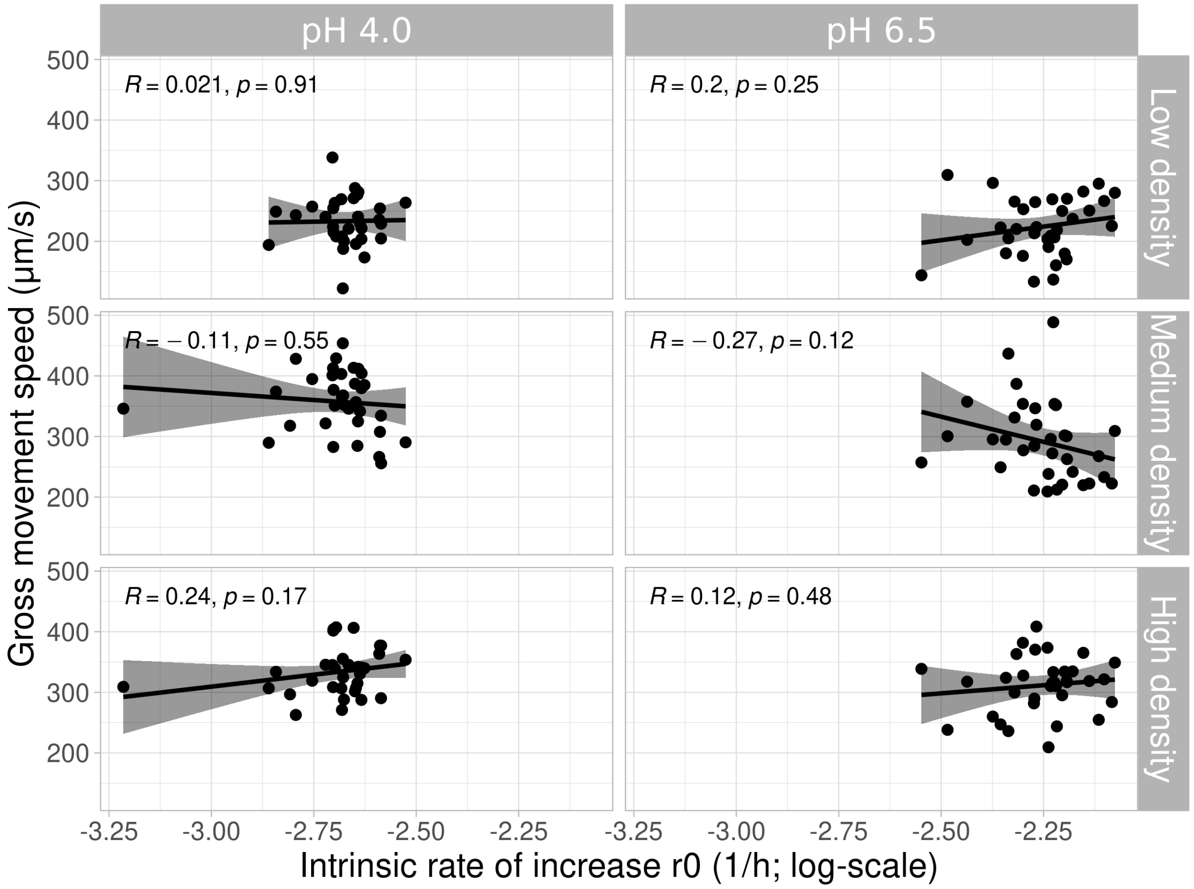


Figure S2: **No correlation between intrinsic growth rate $r_0$ and movement speed.** Pearson correlation between the intrinsic growth rate $r_0$ (x-axis, log-scale) and the gross movement speed (y-axis).

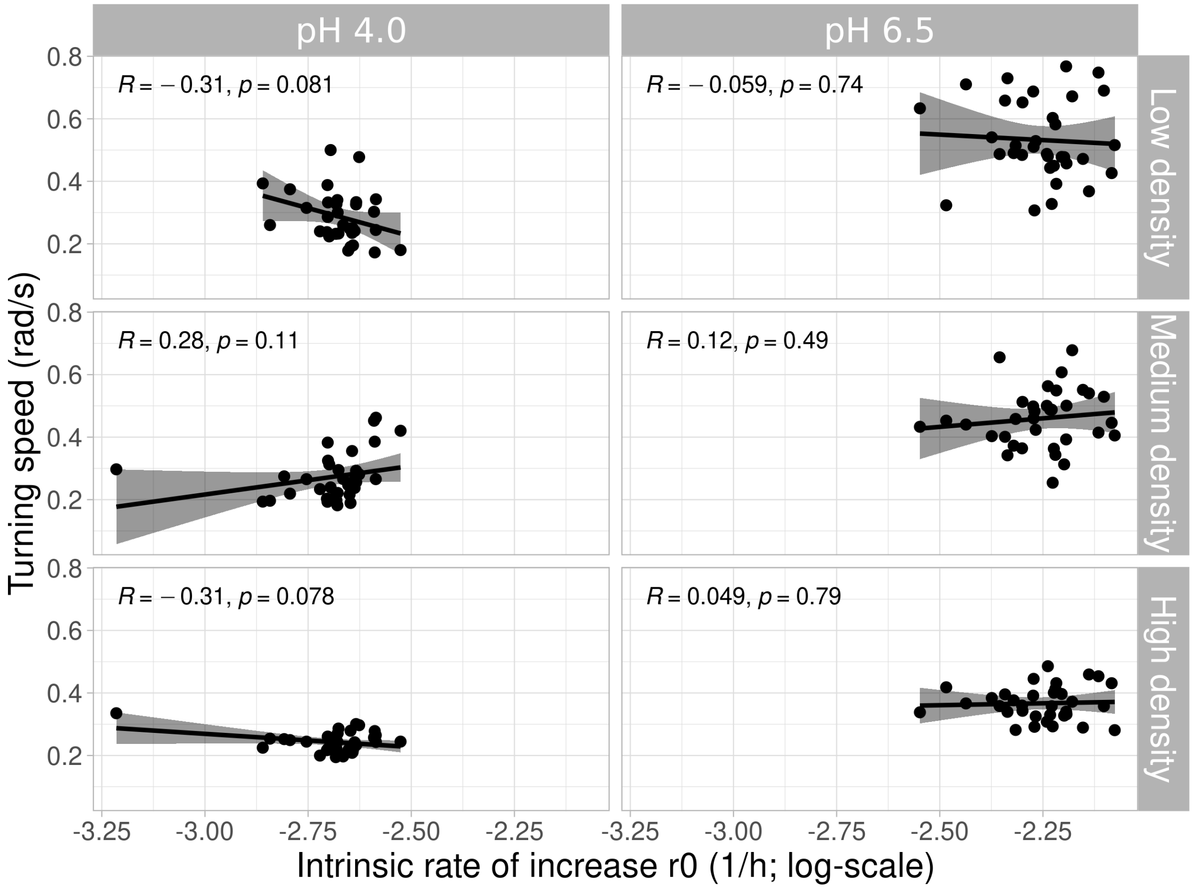


Figure S3: **No correlation between intrinsic growth rate $r_0$ and turning speed.** Pearson correlation between the intrinsic growth rate $r_0$ (x-axis, log-scale) and the turning speed (y-axis).

#### S11 Analysis of fitness change after experimental range expansion

To investigate whether the fitness analysis described in Moerman et al. (2020b) is robust against exclusion of populations that we did not included in our genetic analysis, we repeated this fitness analysis only for those populations we had subjected to pooled genome sequencing. These were the 16 among 34 surviving populations previously evolved by Moerman et al. (2020b) with the highest fitness increase in one of eight experimental conditions (asexual/sexual reproduction, presence/absence of gene flow, presence/absence of a pH-gradient during range expansion, two populations per condition, see also section ”Experimental evolution”). We quantified the change in the intrinsic population growth rate $r_0$ for every evolved population relative to the ancestral population in the pH conditions that those populations experienced during range expansion (i.e., pH 6.5 for populations expanding into a uniform environment, and pH 4.0 for populations expanding into a pH-gradient). Subsequently, we fit a linear model (lm function; R Core Team, 2020) to assess the effect of the mode of reproduction, gene flow, and presence of the pH-gradient (explanatory variables) on the change in $r_0$ (response variable). We then used the dredge function (MuMIn R-package, version 1.43.17; Bartoń, 2009) to find the best fitting model based on the AICc value (Akaike information criterion, corrected for small sample size (Hurvich and Tsai, 1989)). This method allows a ranking of linear models based on the maximum likelihood, where a lower AICc score indicates a better model fit. We report model weights and statistical output (type-III Anova table and model summary) for the best fitting model.

Specifically, this means that we created all models of change in $r_0$ (response variable) as a function of the mode of reproduction, gene flow and presence/absence of a pH-gradient (response variables), ranging from an intercept model to a full interaction model, where all response variables are allowed to interact. We then compared these models using the AICc (Hurvich and Tsai, 1989) to find the best fitting model. This analysis showed that gene swamping (the interaction between reproductive mode and gene flow), and the presence of a pH-gradient affected the fitness change during range expansions.

Specifically, fitness increased in all evolved populations (Figure 2; data from each evolved population is shown as a circle; lines and shaded areas represent model predictions and confidence intervals). Fitness increased more strongly for populations expanding into a pH-gradient (panels B and D in Figure 2) than for populations expanding into a uniform environment (Figure 2 panels A and C; ${\chi^2}_{1,16}$=299.16, p<0.001). This is not surprising, because the low pH experienced during range expansion into a pH-gradient represents a novel environment, whereas the uniform environment more closely matched the abiotic conditions experienced by the ancestor clones.

In addition, we observed a significant interaction between reproduction and gene flow (gene swamping effect; ${\chi^2}_{1,16}$=14.36, p=0.003). Specifically, in the absence of gene flow (Figure 2, panels A and B), the fitness of sexually reproducing populations increased more strongly than that of asexually reproducing populations. In contrast, in the presence of gene flow (Figure 2, panels C and D), sexually reproducing populations increased their fitness less than asexually reproducing populations. This observation suggests that recombination is only beneficial when a population is not being swamped with maladapted genes. We observed this gene swamping effect both in populations expanding into a uniform environment (Figure 2, panels A and C) and in populations expanding into a pH-gradient (Figure 2, panels B and D), although the gene swamping hypothesis only predicts this pattern in the presence of an environmental gradient (Kirkpatrick and Barton, 1997; García-Ramos and Kirkpatrick, 1997). The observations made here are consistent with our previous observations from a larger number of populations (Moerman et al., 2020b).

##### S11.1 Fitness changes: evolution of population growth rate

| | DF | $\chi^2$-value | Pr (>$\chi^2$) |
|---|---|---|---|
| (Intercept) | 1 | 11.42 | 0.006 |
| Reproduction | 1 | 6.44 | 0.027 |
| Gene flow | 1 | 8.09 | 0.016 |
| Abiotic conditions | 1 | 299.16 | <0.001 |
| Reproduction×Gene flow | 1 | 14.36 | 0.003 |
| Residuals | 11 | | |

Table S10 Type III Anova table for the best model describing the change of intrinsic population growth rate, according to AICc model comparisons. For each of the variables, we list,the degrees of freedom (Df) the $\chi^2$ value, and the significance (Pr>$\chi^2$).

| | Estimate | Std. Error | t value | Pr(>\|t\|) |
|---|---|---|---|---|
| (Intercept) | 0.127 | 0.038 | 3.379 | 0.006 |
| Reproduction (Sexual) | 0.121 | 0.048 | 2.539 | 0.028 |
| Gene flow (Present) | 0.136 | 0.048 | 2.844 | 0.016 |

| | | | | |
|---|---|---|---|---|
| Abiotic conditions (Gradient) | 0.583 | 0.034 | 17.296 | <0.001 |
| Reproduction (Sexual)× Gene flow (Present) | -0.256 | 0.067 | -3.789 | 0.003 |

Table S11 Type III Anova table for the best model describing the change of intrinsic population growth rate, according to AICc model comparisons. We list the estimates for the variables (Estimate), the standard error of the estimate (Std. Error), the degrees of freedom (DF), the t-value, and the probability that the estimate is different from 0 (Pr>$|t|$).

### Supplementary Material: Genetic data and results

#### S11.2 Principle component analysis on standing genetic variation

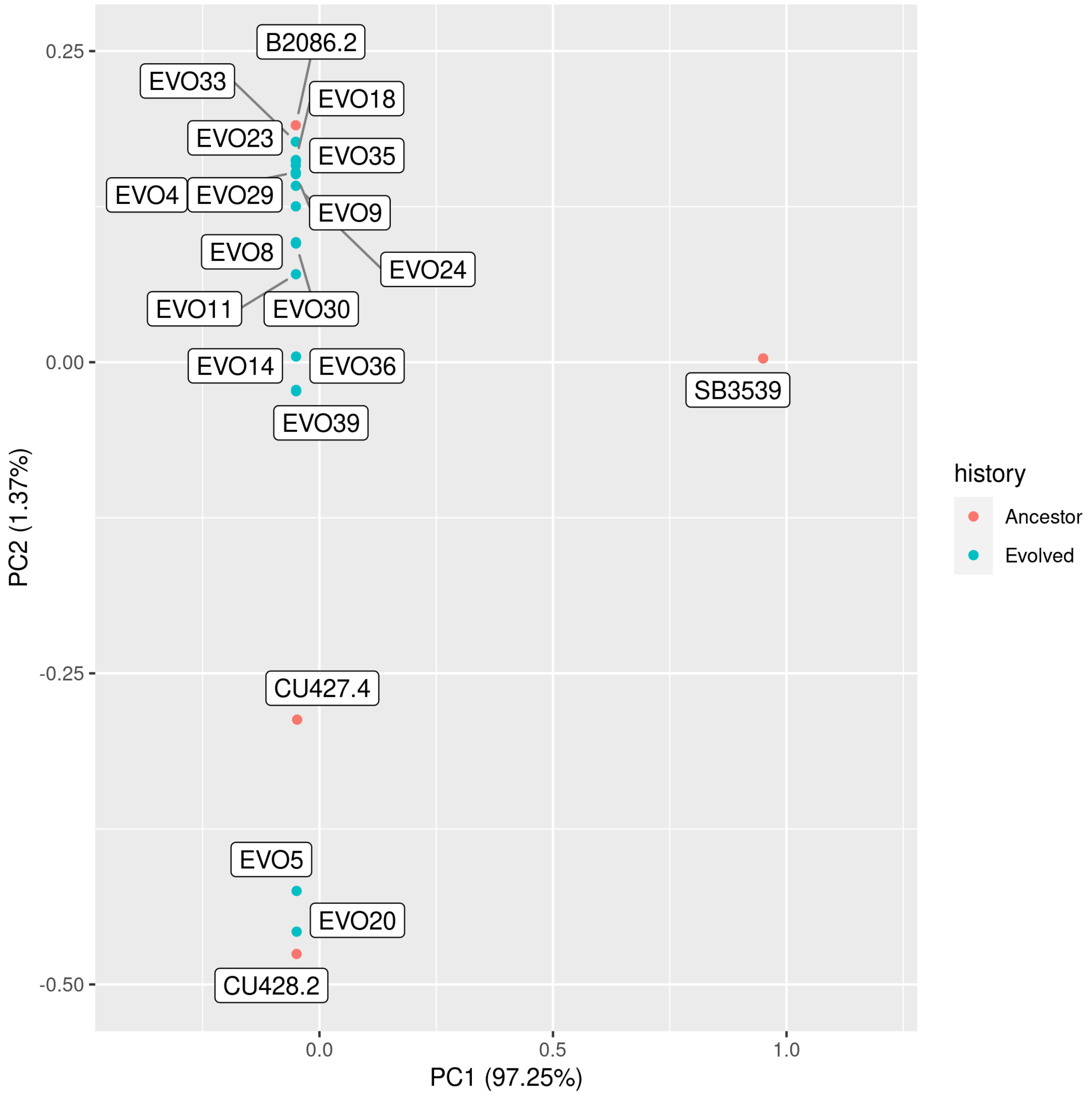


Figure S4: textbfStrong genetic divergence of clone SB3539. Principle component analysis, comparing allele frequencies for standing genetic variation among all 20 populations (4 ancestral clones and 16 evolved populations). Note that almost all variation is associated with PC1 (97.24%), which corresponds to the highly divergent ancestral clone (SB3539).

#### S12 Model summary tables

#### S12.1 The factors that influence changes in standing genetic variation

| | mean | sd | 2.5 % | 97.5 % | n_eff | Rhat4 |
|---|---|---|---|---|---|---|
| Intercept | 8.8499 | 0.1586 | 8.5372 | 9.1585 | 10042 | 1.0000 |
| Cut-off | -9.5299 | 0.2004 | -9.9287 | -9.1384 | 14450 | 0.9999 |
| Reproduction (sexual) | 0.4535 | 0.2111 | 0.0387 | 0.8665 | 10916 | 1.0000 |
| Abiotic conditions (pH-gradient) | 0.3512 | 0.2157 | -0.0678 | 0.7724 | 10417 | 1.0000 |
| Gene flow (present) | 0.6509 | 0.2100 | 0.2359 | 1.0634 | 10731 | 0.9999 |
| Cut-off×reproduction (sexual) | 2.0212 | 0.1967 | 1.6364 | 2.4114 | 15466 | 1.0000 |
| Cut-off×Abiotic condition (pH-gradient) | 2.2032 | 0.2380 | 1.7346 | 2.6705 | 14445 | 1.0000 |
| Cut-off×Gene flow (present) | 0.0838 | 0.1702 | -0.2469 | 0.4184 | 19379 | 1.0001 |
| Reproduction (sexual)×Abiotic conditions (pH-gradient) | 0.1634 | 0.3085 | -0.4436 | 0.7617 | 11884 | 0.9999 |
| Reproduction (sexual)×Gene flow (present) | -0.7596 | 0.2805 | -1.3111 | -0.2086 | 11125 | 0.9999 |
| Abiotic conditions (pH-gradient)×Gene flow (present) | -0.4909 | 0.2963 | -1.0805 | 0.0723 | 11090 | 1.0000 |
| Cut-off×Reproduction (sexual)×Abiotic conditions (pH-gradient) | -5.8767 | 0.3578 | -6.5762 | -5.1863 | 16806 | 0.9999 |
| Cut-off×Reproduction (sexual)×Gene flow (present) | 0.0813 | 1.4003 | -2.6414 | 2.8353 | 21022 | 1.0000 |
| Cut-off×Gene flow (present)×Abiotic conditions (pH-gradient) | -0.9718 | 0.2619 | -1.4902 | -0.4664 | 18851 | 1.0003 |
| Reproduction (sexual)×Abiotic conditions (pH-gradient)×Gene flow (present) | 0.0477 | 1.4027 | -2.6672 | 2.7946 | 20946 | 1.0001 |
| Cut-off×Reproduction (sexual)×Abiotic conditions (pH-gradient)×Gene flow (present) | 4.2403 | 0.3639 | 3.5320 | 4.9464 | 21047 | 1.0000 |

Table S12 **Reproductive mode, gene flow and abiotic conditions affect the incidence of genetic changes from standing variation.** Posterior distributions for the model describing the factors that affect allele frequency changes in standing genetic variation. For each of the variables, we list the mean estimate (log-scale), 2.5 % and 97.5 % quantile of the posterior distribution (log-scale), the estimated effective sample size (n_eff), and the Rhat4 convergence statistic. A large value of n_eff, and a value of Rhat4 close to 1 indicate a good model fit.

#### S12.2 The factors that influence the incidence of *de novo* variants

| | mean | sd | 2.5 % | 97.5 % | n_eff | Rhat4 |
|---|---|---|---|---|---|---|
| Intercept | 1703.0793 | 223.6130 | 1261.6840 | 2140.1009 | 10424 | 1.0000 |
| Cut-off | -1924.7616 | 115.8500 | -2147.1810 | -1695.9409 | 19364 | 1.0000 |
| Reproduction (sexual) | 292.8531 | 278.8231 | -254.6456 | 837.9273 | 14270 | 1.0000 |
| Abiotic conditions (pH-gradient) | -8.2764 | 279.4248 | -560.0772 | 543.1450 | 14371 | 1.0000 |
| Gene flow (present) | 56.2276 | 284.0507 | -499.0587 | 619.6152 | 14506 | 1.0001 |
| Cut-off×reproduction (sexual) | -272.3751 | 134.3106 | -541.0645 | -13.8239 | 21122 | 0.9999 |
| Cut-off×Abiotic condition (pH-gradient) | 34.5827 | 163.2129 | -286.4487 | 353.4795 | 17147 | 1.0000 |
| Cut-off×Gene flow (present) | -74.4128 | 133.5400 | -337.3357 | 185.2668 | 21753 | 0.9999 |
| Reproduction (sexual)×Abiotic conditions (pH-gradient) | 97.2084 | 349.0244 | -596.7746 | 775.4984 | 20290 | 1.0000 |
| Reproduction (sexual)×Gene flow (present) | 29.4462 | 345.5969 | -646.3406 | 698.1330 | 19982 | 1.0001 |
| Abiotic conditions (pH-gradient)×Gene flow (present) | 15.4815 | 347.2358 | -661.0284 | 692.0237 | 18618 | 1.0000 |
| Cut-off×Reproduction (sexual)×Abiotic conditions (pH-gradient) | 92.9039 | 420.2113 | -728.5497 | 914.7278 | 29425 | 0.9999 |
| Cut-off×Reproduction (sexual)×Gene flow (present) | 32.7105 | 211.1267 | -380.8047 | 448.8805 | 19852 | 0.9999 |
| Cut-off×Gene flow (present)×Abiotic conditions (pH-gradient) | 95.4208 | 423.2977 | -731.4223 | 925.3427 | 29083 | 1.0001 |
| Reproduction (sexual)×Abiotic conditions (pH-gradient)×Gene flow (present) | -32.9108 | 239.2737 | -507.1422 | 436.6553 | 22571 | 1.0000 |
| Cut-off×Reproduction (sexual)×Abiotic conditions (pH-gradient)×Gene flow (present) | 162.1835 | 10.5285 | 143.1713 | 184.6253 | 17785 | 0.9999 |

Table S13 **Reproductive mode affects the incidence of *de novo* variants.** Posterior distributions for the model describing the factors that affect the incidence of *de novo* variants. For each of the variables, we list the mean estimate (log-scale), 2.5 % and 97.5 % quantile of the posterior distribution (log-scale), the estimated effective sample size (n_eff), and the Rhat4 convergence statistic. A large value of n_eff, and a value of Rhat4 close to 1 indicate a good model fit.

#### S13 Genetic changes through *de novo* variants and standing genetic variation.

To investigate how all genetic change, both via *de novo* variants and standing genetic variation was affected by the experimental treatments (abiotic conditions, gene flow and reproductive mode), we added the incidence of both types of genetic change for each of the populations. Specifically, this meant we calculated the sum of the number of *de novo* variants and the number of alleles that changed significantly in frequency from standing variation. We then assessed if this total number of genetic variants (i.e. mutations and standing variation) was affected by the mode of reproduction, by gene flow, or by the pH-gradient. In this analysis, we applied various cut-offs (0.3, 0.4, 0.5, 0.6, 0.7 and 0.8) for the magnitude of the allele frequency change. That is, for de novo variants we counted the number of variants whose frequency exceeded this threshold, and for standing variation, we counted the number of alleles whose frequency change in the course of evolution exceeded this threshold . We subsequently fitted a Bayesian generalized linear mixed model with a Poisson distribution to the resulting data, using the Rstan and the "statistical rethinking" package (Stan Development Team, 2020; McElreath, 2015). We created a full interaction model, in which reproduction, gene flow, abiotic conditions, and the cut-off value are all allowed to interact as fixed effects, and in which the identity of the replicate population is included as a random effect. We report posterior distributions (means and 95 % confidence intervals) for the resulting parameter estimates.

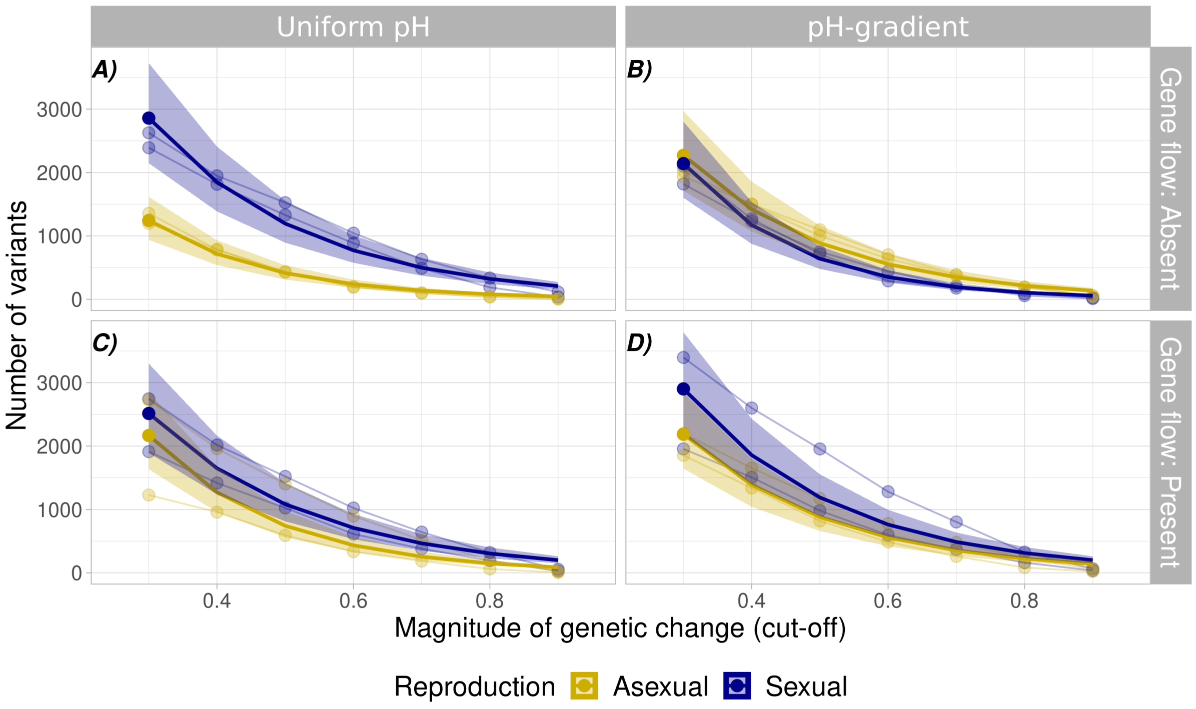


Figure S5: **Mode of reproduction and gene flow alter both the incidence of *de novo* variants and allele frequency changes from standing variation.** The y-axis shows the sum of the incidence of *de novo* variants, and the number of alleles that changed significantly in frequency from standing genetic variation, during range expansion into a uniform environment (panel A and C) or into a pH-gradient (panel B and D). The x-axis shows the cut-of, that is, the minimum frequency that *de novo* variants reached in the population, or the minimum change in allele frequency for variants that changed from standing genetic variation. Circles represent allele frequency data, with thin lines connecting data from the same replicate population. Thick opaque lines show the model predictions for the best model. Shaded areas show 95%-confidence intervals from the posterior distribution. Colours represent mode of reproduction (yellow: asexual, blue:sexual).

We observed similar numbers of variants for populations that expanded into a uniform environment (Figure S5, panels A and C) and into a pH-gradient (Figure S5, panels B and D). We found that sexual reproduction led to more genetic change when populations expanded in a uniform environment and in absence of gene flow (Figure S5, panel A). When either gene flow was present or when populations expanded into a pH-gradient, we did not observe a clear difference between populations with an asexual and sexual mode of reproduction.

| | mean | sd | 2.5 % | 97.5 % | n_eff | Rhat4 |
|---|---|---|---|---|---|---|
| Intercept | 8.8255 | 0.1428 | 8.5481 | 9.1058 | 2377 | 1.0022 |
| Cut-off | -5.6452 | 0.0669 | -5.7770 | -5.5126 | 4133 | 1.0002 |
| Reproduction (sexual) | 0.4758 | 0.1983 | 0.0872 | 0.8638 | 2303 | 1.0018 |
| Abiotic conditions (pH-gradient) | 0.2235 | 0.2013 | -0.1701 | 0.6216 | 2697 | 1.0022 |
| Gene flow (present) | 0.4266 | 0.2001 | 0.0355 | 0.8192 | 2466 | 1.0029 |
| Cut-off×reproduction (sexual) | 1.1785 | 0.0656 | 1.0512 | 1.3079 | 5550 | 1.0001 |
| Cut-off×Abiotic condition (pH-gradient) | 1.1406 | 0.0883 | 0.9687 | 1.3130 | 3888 | 1.0002 |
| Cut-off×Gene flow (present) | 0.3568 | 0.0602 | 0.2404 | 0.4753 | 6189 | 1.0000 |
| Reproduction (sexual)×Abiotic conditions (pH-gradient) | -0.0578 | 0.2804 | -0.6095 | 0.4918 | 2655 | 1.0009 |
| Reproduction (sexual)×Gene flow (present) | -0.6983 | 0.2796 | -1.2385 | -0.1512 | 2477 | 1.0017 |
| Abiotic conditions (pH-gradient)×Gene flow (present) | -0.3398 | 0.2825 | -0.8839 | 0.2056 | 2704 | 1.0016 |
| Cut-off×Reproduction (sexual)×Abiotic conditions (pH-gradient) | -2.7062 | 0.1154 | -2.9335 | -2.4774 | 5153 | 1.0000 |
| Cut-off×Reproduction (sexual)×Gene flow (present) | 0.2310 | 1.4211 | -2.5681 | 3.0086 | 6741 | 1.0000 |
| Cut-off×Gene flow (present)×Abiotic conditions (pH-gradient) | -0.6068 | 0.1020 | -0.8065 | -0.4071 | 5228 | 1.0001 |
| Reproduction (sexual)×Abiotic conditions (pH-gradient)×Gene flow (present) | 0.2193 | 1.4238 | -2.5481 | 3.0209 | 6799 | 0.9999 |
| Cut-off×Reproduction (sexual)×Abiotic conditions (pH-gradient)×Gene flow (present) | 1.8204 | 0.1219 | 1.5795 | 2.0565 | 7430 | 0.9999 |

Table S14 **Reproductive mode, gene flow and abiotic conditions affect the incidence of all genetic changes.** Posterior distributions for the model describing the factors that affect the incidence of *de novo* variants as well as allele frequency changes from standing genetic variation. For each of the variables, we list the mean estimate (log-scale), 2.5 % and 97.5 % quantile of the posterior distribution (log-scale), the estimated effective sample size (n_eff), and the Rhat4 convergence statistic. A large value of n_eff, and a value of Rhat4 close to 1 indicate a good model fit.

#### S14 No association between fitness change and number of *de novo* variants.

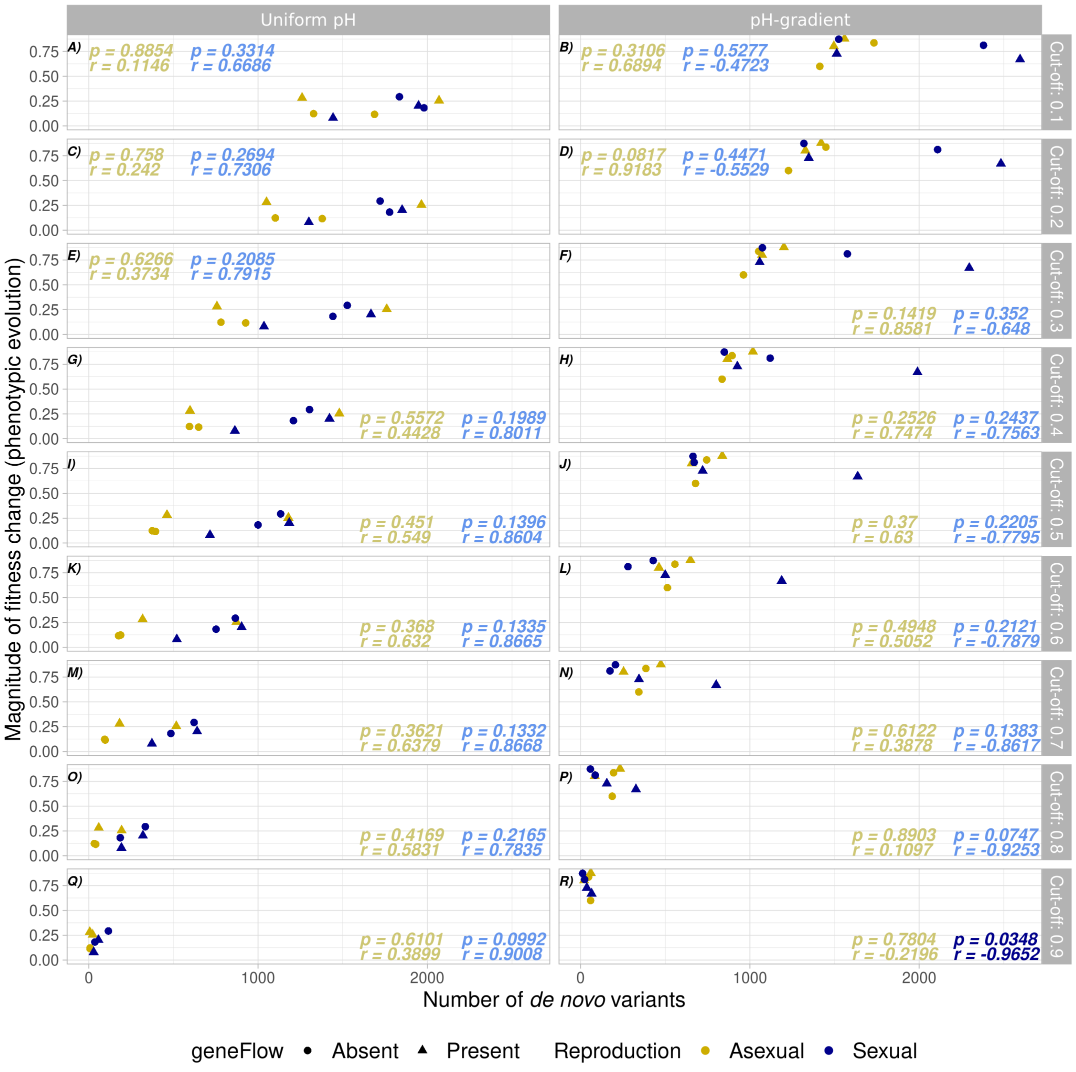


Figure S6: **No association between fitness change and number of *de novo* viariants.** Association between change in fitness and rate of molecular change (number of *de novo* variants). The y-axis shows phenotypic evolution in terms of the log ratio of growth rate change ($log_2(r_0$ of evolved population / $r_0$ of ancestral population)). The x-axis shows the number of *de novo* variants detected in the populations. The left subplots show the data for populations expanding into a uniform environment, and the right subplots for populations expanding into a pH-gradient. Horizontal subplots show the data for different cut-off values for the minimal frequency of *de novo* variants. Each circle within a subplot represents a single population, with colour representing the mode of reproduction (yellow=asexual, blue=sexual) and shape representing gene flow (circle=gene flow absent, triangle=gene flow present). Text insets shows the Pearson correlation coefficient $r$ and significance p, based on a paired sample correlation test for sexual populations (blue text) and asexual populations (yellow text). Non-significant correlations are shown in a faded colour, whereas significant correlations are shown in a brighter colour. Note that we only observed one statistically significant correlation, for sexual populations expanding into a pH-gradient, at the most stringent cut-off (panel R).

#### S15 Genes involved in adaptive evolution through *de novo* variants.

To identify the functions of those genes in which *de novo* variants preferentially occurred, we focused on mutations associated with general adaptations and mutations associated with gradient-specific adaptations.

We considered a *de novo* variant as being associated with a general adaptation if it occurred in at least 75% (12/16) of our evolved populations, independent of reproductive mode, gene flow, or the presence of a pH-gradient.

We considered a mutation to be associated with the presence or absence of a pH-gradient if it had a high prevalence in populations expanding into a pH-gradient and a low prevalence in populations expanding into a uniform environment, or vice versa. To identify such mutations, we first created a list of all genes with at least one *de novo* variant in at least one evolved population. We then scored every evolved population and each of these genes for the presence of mutations. Specifically, for each gene we assigned a value of zero to a population if the gene harboured no mutations in this population, or a value of one if the gene harboured one or more mutations. We then summed these scores separately for the evolved populations expanding into a pH-gradient, and for the populations expanding into a uniform environment to obtain a prevalence of mutations for each class of populations. This prevalence can assume values between zero (no population harbours a mutation in the gene) and eight (all populations do). Finally, we calculated the absolute difference between these prevalences, to obtain a metric for the discrepancy in mutation prevalence between populations that had expanded into a pH-gradient, and populations that had expanded into a uniform environment. We then ranked genes by their difference in this prevalence, and focused on genes for which the absolute difference in prevalence was five or larger. These are genes in which mutations occurred in at least five more populations that had expanded into a pH-gradient than in populations that had expanded into a uniform environment, or vice versa. We discuss the biological functions of these high-discrepancy genes below.

##### S15.1 Analysis of *de novo* variants: general adaptations

We found in total 66 genes with mutations in at least 75% of the evolved populations (12 or more out of 16 populations). Out of these 66 genes, 26 were genes encoding kinase domain proteins or protein kinases. Kinases constitute a fairly large proportion (approximately 3.8%) of the *T. thermophila* proteome (Eisen et al., 2006), yet this alone is not enough to explain the high proportion (39.4%) of kinases in our dataset. Although kinase domain proteins were clearly the most represented group, we found genes in three other functional categories where mutations were common. Specifically, we found common mutations in nine genes encoding transmembrane proteins, in five genes encoding cyclic nucleotide-binding domain proteins, and in two genes encoding translation initiation factors (a full list of all common mutations can be found in Table S18).

##### S15.2 Analysis of *de novo* variants: gradient-specific adaptations

We found a total of 47 genes with large differences in the prevalence of mutations between populations expanding into a gradient and populations expanding into a uniform environment. Out of these 47 genes, genes encoding transmembrane proteins were most common (ten out of 47 genes; 21.28%). Next most common were genes encoding kinase domains (eight out of 47 genes), genes encoding cyclic nucleotide binding proteins (two out of 47 genes) and genes encoding zinc finger proteins (two out of 47 genes). A full list of all gradient-specific mutations can be found in Table S19.
